# Organism-wide secretome mapping uncovers pathways of tissue crosstalk in exercise

**DOI:** 10.1101/2022.11.21.517385

**Authors:** Wei Wei, Nicholas M. Riley, Xuchao Lyu, Xiaotao Shen, Jing Guo, Meng Zhao, Maria Dolores Moya-Garzon, Himanish Basu, Alan Tung, Veronica L. Li, Wentao Huang, Katrin J. Svensson, Michael P. Snyder, Carolyn R. Bertozzi, Jonathan Z. Long

## Abstract

There has been growing interest in identifying blood-borne factors that mediate tissue crosstalk and function as molecular effectors of physical activity. Although past studies have focused on an individual molecule or cell type, the organism-wide secretome response to physical activity has not been evaluated. Here, we use a cell type-specific proteomic approach to generate a 21-cell type, 10-tissue map of exercise-regulated secretomes in mice. Our dataset identifies >200 exercise-regulated cell type-secreted protein pairs, the majority of which have not been previously reported. *Pdgfra*-cre-labeled secretomes were the most responsive to exercise training. Elevated lactate levels can directly regulate protein secretion. Lastly, we establish an anti-obesity and anti-diabetic role for a proteoform of an intracellular carboxylesterase whose secretion from the liver is induced by exercise training. Together, our data uncover the dynamic remodeling of cell and tissue crosstalk by physical activity.

## INTRODUCTION

Physical activity is a powerful physiologic stimulus that provides benefits to many organ systems and confers protection against disease (Piercy et al., 2018; Warburton and Bredin, 2017). Conversely, physical inactivity is a major contributor to cardiovascular morbidity and mortality (Booth et al., 2017; Lear et al., 2017). The magnitude of the benefits of physical activity is comparable, and in some cases even greater, than currently available first-line pharmacological treatments (Blair et al., 1989; Hambrecht et al., 2004; Knowler et al., 2002). The mechanisms responsible for the benefits of exercise are incompletely understood, but likely extend beyond activity-associated increases in energy expenditure alone (McGee and Hargreaves, 2020; Neufer et al., 2015).

In recent years, there has been tremendous interest in the identification and characterization of exercise-inducible, soluble (e.g., secreted) blood-borne molecules. These circulating molecules, which have been called “exerkines” or “exercise factors,” are secreted signaling molecules that function as molecular effectors of physical activity (Chow et al., 2022; Safdar et al., 2016). Over 50 years ago, Goldstein (Goldstein, 1961) demonstrated that contracting muscle from dogs produced a humoral factor that stimulated glucose uptake when transferred to non-exercised muscle preparations. More recently, experiments involving re-infusion of exercise-conditioned plasma in mice have also provided additional evidence for bioactive molecules present in the circulation following exercise (De Miguel et al., 2021; Horowitz et al., 2020).

At a molecular level, many individual candidate metabolites, lipids, polypeptides and proteins from diverse tissues have been proposed to function as exerkines (Agudelo et al., 2014; Boström et al., 2012; Knudsen et al., 2020; Lynes et al., 2017; Rao et al., 2014; Reddy et al., 2020; Sato et al., 2022; Steensberg et al., 2000; Takahashi et al., 2019; Wrann et al., 2013; Parker et al., 2017). However, these previous efforts have typically focused on a single factor (e.g., IL-6) and/or a single cell type/tissue of origin (e.g., muscle). Few studies have systematically mapped exercise-inducible secreted molecules across an entire organism. A major challenge, especially for secreted polypeptides and proteins, has been the low depth of plasma proteome coverage by classical shotgun proteomics techniques (Uhlén et al., 2015). Aptamer- and antibody-based approaches provide higher sensitivity, but are not comprehensive for the plasma proteome and cannot be used to detect the array of potentially new proteoforms or cleavage fragments that might be produced following physical activity (Boström et al., 2012; Somineni et al., 2014). Finally, many secreted proteins are expressed by multiple cell types, and single snapshot detection of these molecules in the circulation would not be expected to enable detection of cell type-specific, and potentially bidirectional regulation of exercise-inducible changes across distinct cell types.

We (Wei et al., 2021; Wei et al., 2020) and others (Droujinine et al., 2021; Kim et al., 2021; Liu et al., 2021) have recently described a biochemical secretome profiling methodology that enables direct labeling, enrichment, and identification of secreted proteins in mice at a cell type-specific resolution. Key to this methodology is the delivery of an engineered biotinylation enzyme TurboID (Branon et al., 2018) into the secretory pathway of cells via adeno-associated virus (AAV) transduction. Cell type-specific labeling is achieved genetically because the expression of the TurboID is restricted to those cells expressing cre recombinase. Biotinylated and secreted plasma proteins can then be purified directly from blood plasma using streptavidin beads and analyzed by LC-MS/MS. Our initial secretome studies with the proximity biotinylation approach previously established that cell type-specific secretome changes could be profiled directly in an intact animal (Wei et al., 2020). Here, we have dramatically extended this secretome profiling to organism-wide scale and examined the cell type-specific secretome response to exercise training. Our organism-wide 21-cell type, 10-tissue secretome map of physical activity provides fundamental insights into the molecular identity and cellular origin of exercise-regulated circulating factors and illuminates the dynamic regulation of intercellular and inter-organ crosstalk by physical activity.

## RESULTS

### Study design and proteomic map of exercise-regulated secretomes

The experimental design for organism-wide, cell type-specific secretome profiling in response to exercise is shown schematically in **Fig. 1A**. First, we generated adeno-associated adenovirus serotype 9 (AAV9) expressing a cre-inducible, endoplasmic reticulum-restricted TurboID (AAV9-FLEx-ER-TurboID, 3*10e11 GC/mouse, intravenously) (**Fig. 1A** and see **Methods**). In cells expressing cre recombinase, inversion of the FLEx cassette results in robust expression of the ER-TurboID construct. AAV9 was chosen since this serotype exhibits broad tissue distribution and transduction (Zincarelli et al., 2008). Next, we assembled 21 manually-curated hemizygous cre mouse driver lines from the Jackson Laboratories (N = 6/genotype, **Fig. 1A**, see **Methods**). Included amongst the set of cre driver lines were those that target specific organs previously shown to participate in response to exercise (e.g., *MCK-*cre targeting muscle, *Adiponectin-*cre targeting fat, and *Albumin-*cre targeting liver). Other cre drivers exhibited well-validated expression patterns (e.g., *Lysm*-cre for macrophages), even though those cell types had not been previously implicated in the response to physical activity. In the cases where tamoxifen-inducible cre driver lines were used, tamoxifen (2 mg/mouse, intraperitoneally) was administered two weeks after viral transduction. As expected, robust *TurboID* mRNA expression was detected in tissues from all transduced mice compared to virally-transduced, wild-type control mice (**Fig. S1A** and see **Methods**). Furthermore, we harvested a collection of tissues from a subset of cre driver mice in which TurboID expression was predicted to occur in an organ-restricted manner (*Albumin-*cre for liver, *Ucp1*-cre for brown fat, *Adiponectin*-cre for all adipose, *MCK*-cre for muscle, *Myh6*-cre for heart, *Pdx1*-cre for pancreas, and *Syn1*-cre for brain). V5-tagged TurboID protein, as measured by Western blot using an anti-V5 epitope tag antibody, was robustly detected with the expected organ-restricted pattern (**Fig. S1B**).

**Fig 1.**
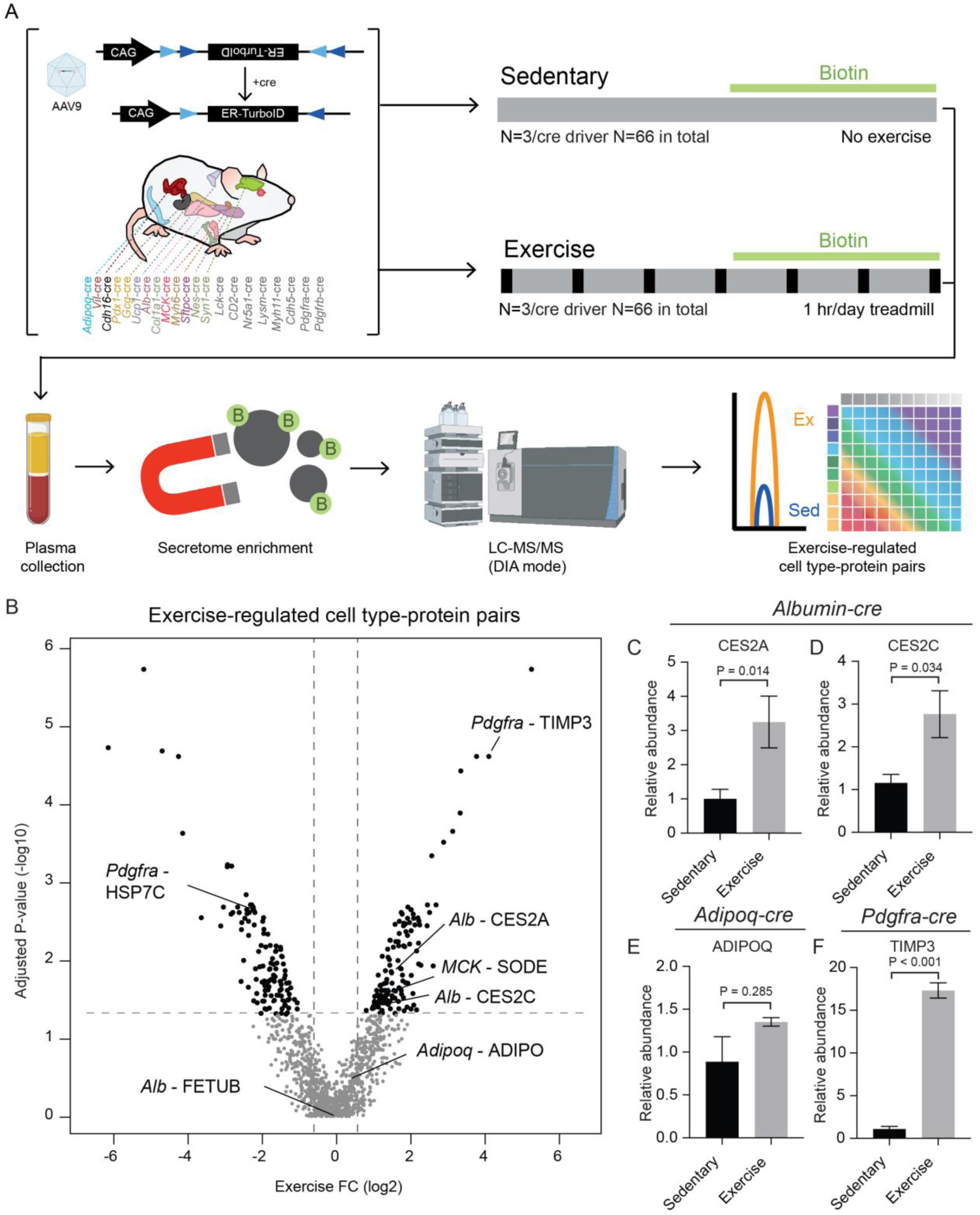
Study design and overview of exercise secretomes across 21 cell types in mice. (A) Overview of the study design including viral transduction (AAV9-FLEx-ER-TurboID, 3*10e11 GC/mouse, intravenously) of 21 cre driver lines (male, N = 3/condition/genotype, see Methods) and C57BL/6 mice (male, N = 3/condition), 1-week treadmill running (20 m/min for 60min per day), secretome labeling (biotin delivered via biotin water (0.5 mg/ml) and via injection (24 mg biotin/ml, intraperitoneally, in a solution of 18:1:1 saline:Kolliphor EL:DMSO, final volume of 200 µl per mouse per day) in the last three days of running), enrichment of biotinylated plasma proteins using streptavidin beads and proteomic analysis. (B) Volcano plot of adjusted P-values (- log10) and exercise fold change (log2) of total 1272 cell type-protein pairs. Adjusted P-values were calculated from moderated t-statistics (see **Methods**). Black dots indicate exercise-regulated cell type-protein pairs (adjusted P-values < 0.05) and gray dots indicate unchanged cell type-protein pairs (adjusted P-values > 0.05). (C-F) Relative abundance of exercise-regulated (C, D and F) and exercise-unregulated (E) cell type-protein pairs from exercise and sedentary mice. N = 3/genotype/condition. In (C-F), P-values were calculated from two-tailed unpaired t-tests.

Three weeks after viral transduction, mice were separated into treadmill running or sedentary groups (N = 3 per group per genotype, **Fig. 1A**). Our one-week treadmill running protocol, adapted from (Wu et al., 2011), consisted of running for 60 min/day at a speed of 20 m/min (see **Methods**). Sedentary mice were kept in their home cages. After the one-week treadmill training protocol, mRNA levels of *Pgc1a* and *Nr4a1* in quadriceps muscle were induced by 2.7-fold (P < 0.05) and 1.6-fold (P < 0.05), respectively (**Fig. S1C**) (Finck and Kelly, 2006; Kawasaki et al., 2009). Total inguinal white adipose tissue mass was reduced by 30% in the exercise group (sedentary: 333 ± 24 mg; exercise: 235 ± 16 mg, mean ± SEM, P < 0.05) (**Fig. S1D**). Histological analysis of the inguinal adipose tissue also showed reduced adipocyte size (**Fig. S1E**). Additionally, all exercised animals exhibited -0.68 ± 0.02 g (mean ± SEM, N = 66) weight change during the 1-week treadmill running whereas the sedentary controls (N = 66) gained +0.50 ± 0.02 g (mean ± SEM, N = 66, **Fig. S1F**). Taken together, these molecular and physiologic data validate both the secretome labeling mouse lines as well as the one-week exercise training protocol.

To determine the identity of exercise-regulated secreted proteins and their cell types of origin, we supplemented biotin to mice for the final three days of the exercise training protocol to biotinylate in vivo secretomes (**Fig. 1A**, right, and see **Methods**). Two hours after the final bout of running, blood plasma was collected from each mouse and biotinylated secreted proteins were purified using streptavidin beads, digested following an S-trap protocol, and analyzed by LC-MS/MS in data-independent acquisition (DIA) mode (see **Methods**). We chose to use a previously described spectrum library-free DIA approach that relies on gas-phase fractionation (GPF)-DIA data from a pooled sample to generate DIA-only chromatogram libraries (Pino et al., 2020a; Searle et al., 2018). This allowed us to search all data against experiment-specific chromatogram libraries using the freely available EncyclopeDIA platform (Searle et al., 2018), followed by further processing in the Skyline and Perseus data analysis environments (Pino et al., 2020b; Tyanova et al., 2016). To filter for bona fide cell type-protein pairs enriched by streptavidin, we also applied a 1.5-fold enrichment filter for each cell type-protein pair versus non-transduced, wild-type controls (see **Methods**). In total across all samples (N = 3 mice/condition x 2 conditions x 21 genotypes), we detected 1,272 unique cell type-protein pairs with ≥ 2 peptides detected in all 3 replicates of both conditions (**Table S1**).

Exercise-regulated cell type-protein pairs were identified by comparison of differential secreted proteins from the same cell type in sedentary versus exercised mice. This comparison also provides a natural control for differing levels of proximity labeling enzyme expression and/or varying levels of secretome biotinylation between distinct cell types. Exercise significantly altered 256 cell type-protein pairs (20.1% of the entire dataset, adjusted P-value < 0.05, **Fig. 1B**), many of which have not been previously described. We identified several example candidates of exercise-regulated secreted proteins selectively altered in one cell type (**Fig. 1C-F**). For instance, two carboxylesterases, CES2A and CES2C, were increased by 3-fold exclusively in secretomes from *Albumin-*cre mice following treadmill running (**Fig. 1C, D**). CES2 enzymes are classically annotated as intracellular, liver-enriched endoplasmic reticulum resident proteins. Nevertheless, our data suggest that CES2 enzymes can also be released from the liver in an exercise-dependent manner. Other well established secreted protein-cell type pairs, such as the hormone adiponectin from fat cells (ADIPOQ in *Adipoq-*cre transduced secretomes) (Stern et al., 2016), was robustly detected in our dataset but not regulated by physical activity (**Fig. 1E**). Finally, one of the most exercise-inducible cell type-protein pairs in the dataset was TIMP3 from *Pdgfra*-cre labeled secretomes (**Fig. 1F**). While TIMP3 had previously been implicated in diverse physiologic roles including in myogenesis (Liu et al., 2010), thermogenesis and metabolism (Hanaoka et al., 2014), vascular remodeling (Basu et al., 2013), and atherogenesis (Stöhr et al., 2014), its exercise-inducible secretion from *Pdgfra*-cre labeled cells has not been previously identified. We conclude that one week of treadmill running in mice results in specific modulation of a subset of secreted proteins-cell type pairs across our entire secretome dataset.

### Systematic analysis of exercise-regulated cell type-secreted protein pairs

The distribution of these exercise-regulated cell type-protein pairs across the 21 cell types is shown in **Fig. 2A**. We observed an average and median of 12 and 10, respectively, proteins changed per cell type secretome in response to exercise, with a range across all cell types of 1 to 44. The number of proteins changed in each cell type secretome by exercise was not simply correlated to secretome size since the two cell types with the largest number of unique proteins identified in their secretomes (*Albumin*-cre, with 142 proteins, and *MCK*-cre, with 132 proteins) exhibited near the median number of exercise-regulated secreted protein changes (**Fig. 2A**). All cell types exhibited some secretome changes following exercise, demonstrating that all cell types exhibit exercise responsiveness to some degree as measured by changes to their production of soluble factors. Approximately half of the exercise-regulated secretome changes (50%, 129 out of 256) were observed in only one cell type (**Fig. 2B**). In addition, the frequency of either exercise up-or down-regulated proteins was equivalent (66 proteins increased and 63 proteins decreased, **Fig. 2B**), suggesting that exercise regulation of soluble factors does not only involve production of new secreted molecules, but also suppression and other regulation of active protein secretion. The other half of the exercise-regulated secretome changes (50%, 127 out of 256) were proteins expressed in multiple cell types that exhibited cell type-specific regulation following exercise. In this latter group, bidirectional change, defined as up-regulation in one cell type and down-regulation in a different cell type, was commonly observed (40%). These data further underscore the increased resolution afforded by cell type-specific secretome profiling, as well as the need to use global approaches for evaluating exercise-induced changes, which are complex, bidirectional, and cell type-specific.

**Fig 2.**
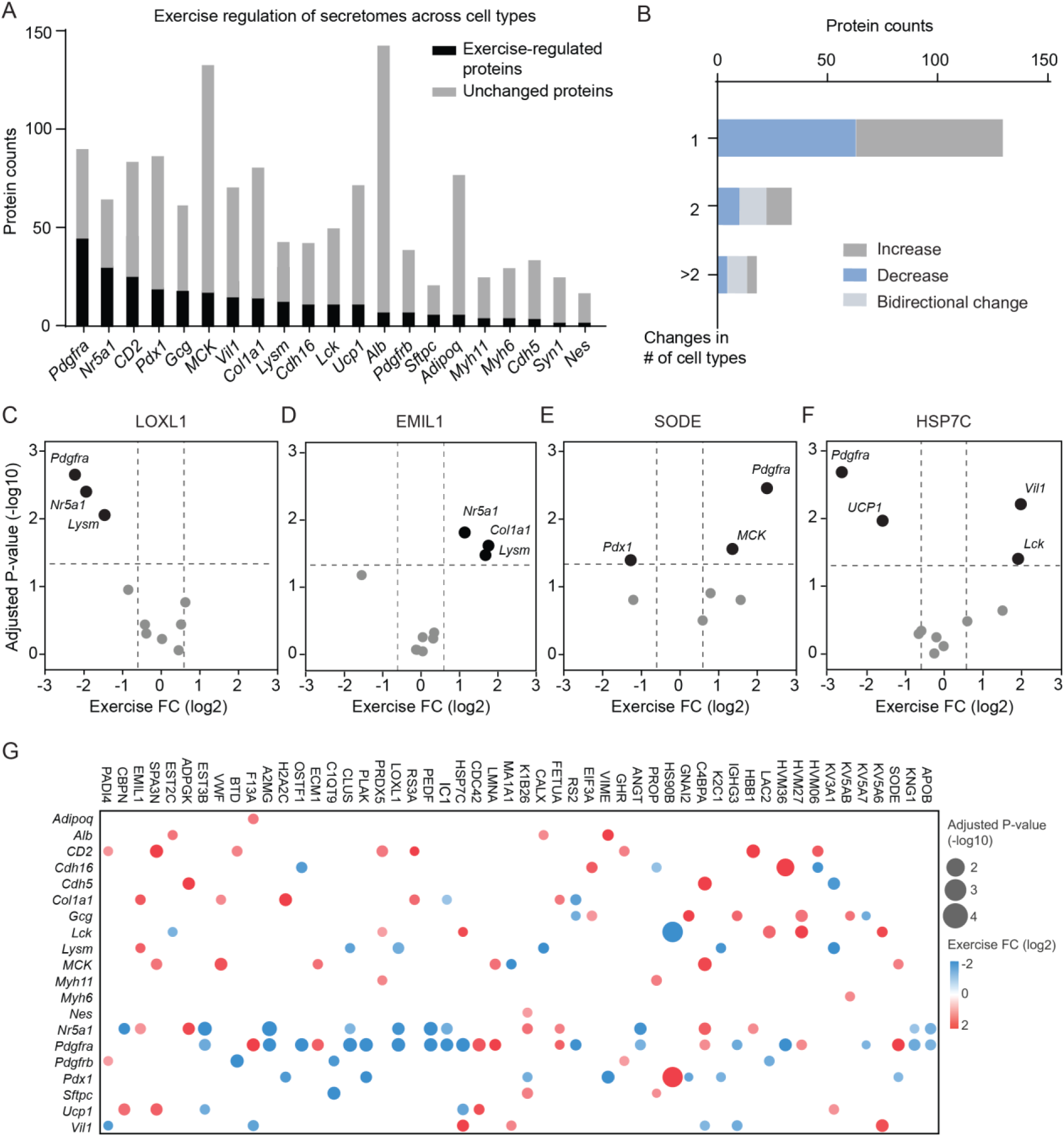
Systematic analysis of exercise-regulated cell type-protein pairs. (A) Bar graph of exercise-regulated proteins (black) and unchanged proteins (gray) across 21 cell types. (B) Histogram of increased (gray), decreased (blue) and bidirectionally changed (light gray) secreted proteins after exercise. (C-F) Volcano plot of adjusted P-values (-log10) and exercise fold change (log2) of indicated example proteins. Black dots indicate exercise-regulated cell type-protein pairs (adjusted P-values < 0.05) and gray dots indicate unchanged cell type-protein pairs (adjusted P-values > 0.05). (G) Bubble plot of adjusted P-values (-log10) and exercise fold change (log2) of proteins changed in more than 1 cell type after exercise. Red dots indicate increased proteins after exercise and blue dots indicated decreased proteins.

Direct examination of the exercise-regulated secreted protein changes of proteins that were expressed across multiple also revealed previously unknown and unusual cell type-specific patterns of exercise regulation (**Fig. 2C-F**). For instance, LOXL1, a secreted enzyme involved in extracellular protein lysine oxidation, was selectively downregulated following exercise in secretomes from *Pdgfra-*, *Nr5a1-*, and *Lysm*-cre transduced mice (**Fig. 2C**). Conversely, the extracellular matrix protein EMIL1 was upregulated following exercise selectively in *Col1a1*, *Lysm* and *Nr5a1*-cre secretomes (**Fig. 2D**). We also identified several examples of proteins expressed in multiple cell types and bidirectionally regulated by exercise. These include SOD3 (upregulated in *MCK*-cre and *Pdgfra*-cre, and downregulated in *Pdx1*-cre secretomes) (**Fig. 2E**) and HSP7C (upregulated in *Vil1*-cre and *Lck*-cre, and downregulated in *Pdgfra*-cre and *UCP1*-cre secretomes) (**Fig. 2F**). A bubble plot visualizing all such examples of exercise-regulated secreted proteins with regulation in ≥ 2 cell type secretomes is shown in **Fig. 2G**. While several of these proteins have been identified as exercise-regulated proteins in the literature, our dataset further contextualizes and refines their interpretations at a cell type level. For instance, total plasma SOD3 was previously reported to be induced by acute treadmill running in humans as well as voluntary wheel running and treadmill exercise in mice (Abdelsaid et al., 2022; Fukai et al., 2000; Hitomi et al., 2008). However, only one of these previous studies suggested that muscle is a cellular origin for exercise-inducible SOD3 (Hitomi et al., 2008). Our dataset not only confirms upregulation of SOD3 secretion from muscle following exercise, but also identifies *Pdgfra* and *Pdx1* as additional cell types that express SOD3 and contribute to the exercise-inducible regulation of this protein (**Fig. 2E**). Similarly, PEDF is a neurotrophic growth factor widely expressed across multiple cell types. In humans, circulating PEDF is reduced after a single bout of cycling (Raschke et al., 2013) as well as after 12-month moderate-intensity aerobic exercise (Duggan et al., 2014). PEDF was downregulated in our dataset in both *Nr5a1*-cre and *Pdgfra*-cre secretomes (**Fig. 2G**), suggesting that these two cell types contribute to the downregulation of plasma PEDF after exercise.

### Secretomes from *Pdgfra-*cre labeled cells are highly responsive to exercise training

While metabolic cell types (muscle, liver, and fat) have typically been examined in the context of exercise, our dataset indicated that *Pdgfra*-cre labeled secretomes were amongst the most exercise-responsive, as measured by the number of exercise-regulated proteins (**Fig. 2A**). As an alternative method for determining magnitude of the exercise response, we also developed a direct “exercise-responsiveness” metric for each cell type secretome in our dataset (see **Methods**). This metric takes into account magnitude and statistical significance of the exercise-regulated changes, as well as the secretome size of that cell type. *Pdgfra*-cre labeled secretomes once again emerged as the most responsive cell type (exercise-responsiveness score = 115.4 and **Fig. 3A**).

**Fig 3.**
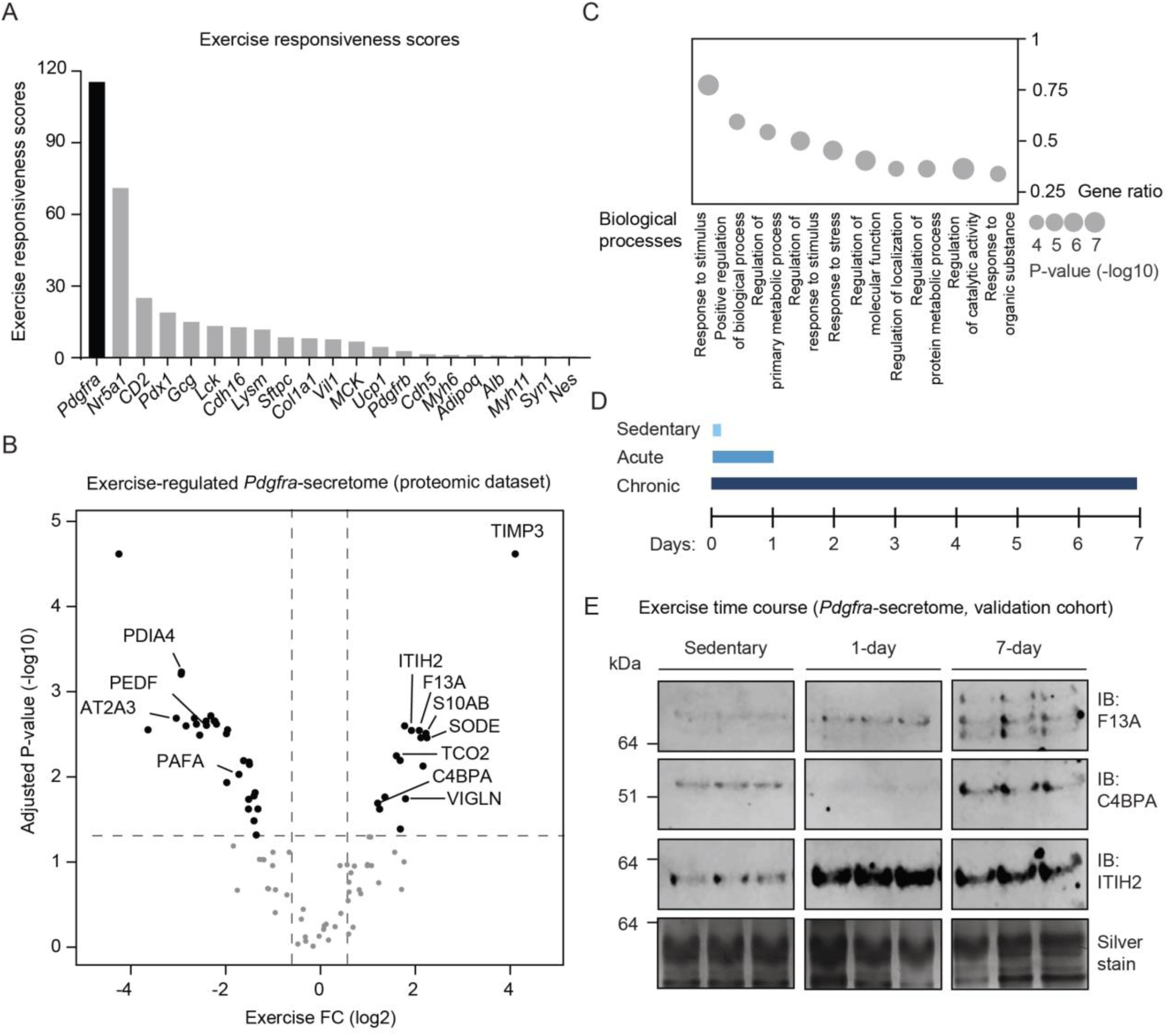
Characterizations of exercise secretomes from *Pdgfra*-cre labeled cells. (A) Bar graph of exercise responsiveness scores across cell types. Exercise responsiveness scores of a given cell type were calculated by summarization of the score of individual exercise regulated protein (adjusted P-values < 0.05) of that cell type with the following equation: sum(absolute exercise fold change (log2) x confidence of the change (-log10(adjusted P-values))) x percent of secretome change (number of exercise regulated proteins (adjusted P-values < 0.05) / total number of secreted proteins of that cell type). See **Methods**. (B) Volcano plot of adjusted P-values (-log10) and exercise fold change (log2) of *Pdgfra* secretomes. Black dots indicate exercise-regulated cell type-protein pairs (adjusted P-values < 0.05) and gray dots indicate unchanged cell type-protein pairs (adjusted P-values > 0.05). (C) Gene ontology analysis of exercise regulated proteins (adjusted P-values < 0.05) from *Pdgfra* secretomes. Size of bubbles represents P-values (-log10) of biological processes enrichment and y axis represents gene ratio. (D) Study design of secretome analysis of heterozygous *Pdgfra*-cre mice (12-week-old male, N = 3/condition) injected with 3*10e11 GC/mouse AAV9-FLEx-ER-TurboID and tamoxifen. Three weeks after tamoxifen delivery, these mice were subjected to acute running (single bout, 20 m/min for 60 min), 1-week chronic running (daily, 20 m/min for 60 min) or being sedentary. Secretome labeling was initiated via injection (24 mg biotin/ml, intraperitoneally, in a solution of 18:1:1 saline:Kolliphor EL:DMSO, final volume of 400 µl per mouse per day) in the last bout of running and biotinylated plasma proteins were enriched using streptavidin beads and analyzed by western blotting (see **Methods**). (E) Anti-F13A (top), anti-C4BPA (second row), anti-ITIH2 (third row) of eluted biotinylated plasma proteins from streptavidin beads after immune purification. Silver stain of total eluted biotinylated plasma proteins was used as loading control (bottom row). Samples (N = 3/condition) were from the experiment described in the legend of Fig. 3D.

*Pdgfra*-cre labeled cells are a population of anatomically-distributed cells that have been described in the literature as fibroblasts, mesenchymal stem cells, or progenitor/precursor cells (Li et al., 2018; Merrick et al., 2019; Zepp et al., 2017). They have diverse roles in tissue remodeling, fibrosis and cell proliferation depending on the resident tissue and physiological context (Li et al., 2018; Zepp et al., 2017). To understand the organ localization of the *Pdgfra*-cre labeled cells labeled in our secretome labeling experiments, we measured *TurboID* mRNA across multiple tissues from *Pdgfra*-cre transduced mice (**Fig. S2A**). Robust *TurboID* mRNA enrichment was detected across many tissues examined, including lung, adipose tissues (inguinal and brown), muscle, gut, kidney, and brain (**Fig. S2A**). This distribution is similar to the reported expression of *Pdgfra* mRNA across tissues and suggest that *Pdgfra* localized to multiple organs are participating in the exercise-regulated secretome response detected in our dataset (**Fig. S2B**).

The entire secretome of *Pdgfra*-cre labeled cells is shown in **Fig. 3B**. A diverse array of secreted proteins with pleiotropic physiologic functions were found to be regulated by exercise. Some of the most upregulated molecules included the previously mentioned TIMP3 (17-fold) and SOD3 (5-fold), as well as the S100 family member S10AB (5-fold), the HDL-binding protein VIGLN (4-fold), and the vitamin B12 transport protein TCO2 (3-fold). Conversely, downregulated secreted proteins in the *Pdgfra*-cre secretome included ER-resident proteins such as the protein disulfide isomerase PDIA4 (87% suppression) and ER calcium ATPase AT2A3 (88% suppression), the growth factor PEDF (81% suppression), and the lipid metabolizing enzyme PAFA (69% down). Gene ontology analysis revealed enrichment of several biological pathways in the exercise-regulated *Pdgfra*-cre secretome, of which the highest scoring by gene ratio was “response to stimulus” (P = 4.96E-07, gene ratio = 0.77, **Fig. 3C**). Additional biological processes identified from the *Pdgfra*-cre secretome included “response to stress” (P = 2.00E-06, gene ratio = 0.45), and “response to organic substance” (P = 1.03E-04, gene ratio = 0.34) (**Fig. 3C**). These observations suggest that *Pdgfra*-cre labeled cells respond to exercise by sensing exercise-regulated environmental cues, such as metabolites, cytokines, or other signaling molecules, that in turn drive bidirectional changes in the *Pdgfra*-cre labeled secretome.

We next sought to validate some of the *Pdgfra*-cre secretome responses using an orthogonal method, as well as to determine whether the secretome changes of *Pdgfra*-labeled cells represented an acute or chronic response to physical activity. Towards this end, we used Western blotting with commercially available antibodies to determine the levels of three exercise-regulated secreted proteins (F13A, C4BPA and ITIH2) from the *Pdgfra-*cre secretome in a new cohort of virus-transduced, *Pdgfra*-cre mice. In our original proteomic dataset, F13A, C4BPA, and ITIH2 were found to be 4-, 2-, and 4-fold upregulated in *Pdgfra*-cre secretomes after 1 week treadmill running. In this experiment, transduced animals (N = 5/group) were separated into three groups: sedentary, acute exercise (single treadmill bout, 20 m/min for 60min), or 1-week chronic exercise (daily treadmill running, 20 m/min for 60 min) (**Fig. 3D**). As shown in **Fig. 3E**, Western blotting revealed that two of the three proteins, F13A and ITIH2, were elevated from *Pdgfra*-cre secretomes in both the acute as well as chronic treadmill running cohorts. Interestingly, while ITIH2 exhibited a similar magnitude upregulation after 1- or 7-days running, the upregulation of F13A was higher as the exercise duration increased (e.g., 7-day > 1 day > sedentary). On the other hand, C4BPA was down-regulated at 1 day, and upregulated by 7-days, indicating this protein is suppressed after acute exercise but increased after chronic training. Importantly, the total eluted proteins from streptavidin purification remained constant (**Fig. 3E**). These data demonstrate that the exercise-regulated proteins in the *Pdgfra*-cre secretome are robust across multiple cohorts and reflect both acute as well as chronic aspects of exercise training.

### In vitro studies link lactate levels to protein secretion

Two of the most robust exercise-inducible molecules from *Albumin-*cre secretomes belonged to the same family of carboxylesterase enzymes (CES2A and CES2C, **Fig. 4A**). Because hepatocytes can be easily cultured and manipulated in vitro, we used the *Albumin-*cre/CES2 cell type-protein pair as a representative molecular handle to investigate the molecular drivers of this process in vitro.

**Fig 4.**
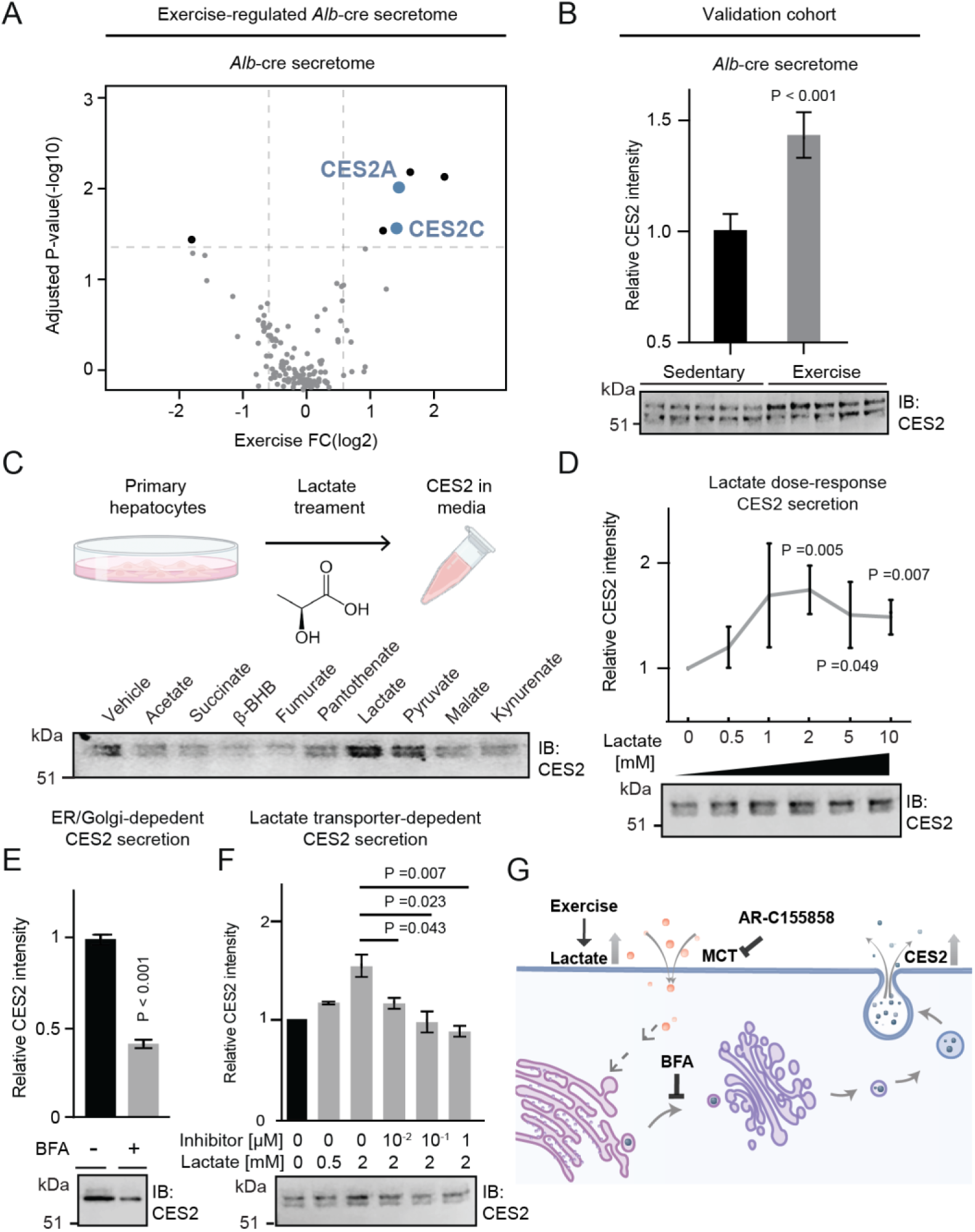
Lactate-induced CES2 secretion in mouse primary hepatocytes. (A) Volcano plot of adjusted P-values (-log10) and exercise fold change (log2) of Albumin secretomes. Black dots indicate exercise-regulated cell type-protein pairs (adjusted P-values < 0.05) and gray dots indicate unchanged cell type-protein pairs (adjusted P-values > 0.05). (B) Anti-CES2 (bottom) blotting and quantifications of band intensity (top) of immune purified biotinylated plasma proteins from 10-week-old Albumin-cre male mice transduced with 3*10e11 vg AAV9-FLEx-ER-TurboID virus and exercised on a treadmill for 1-week. N = 5/group. (C-F) Anti-CES2 blotting (bottom) and quantifications of band intensity (D-F) of conditioned medium of primary hepatocytes isolated from 8 to 12-week-old male C57BL/6J mice. Cells were treated with 2 mM indicated organic compounds (C), 2 mM sodium lactate (D), sodium lactate (2 mM) and BFA (5 μg/ml) (E), indicated concentrations of sodium lactate and AR-C155858 (F) for 4 h before analysis. Experiments in each panel were repeated four times and similar results were obtained. (G) Model of CES2 secretion from cells. Exercise-inducible rise of extracellular lactate induces release of ER-lumen-resident CES2 from hepatocytes. Functional lactate transporters and ER-Golgi vesicle transport are required for CES2 secretion. P-values for quantifications in this figure were calculated from two-tailed unpaired t-tests.

To first validate the exercise-inducible secretion of CES2 proteins from the liver, we used a commercially available pan anti-CES2 antibody to probe streptavidin-purified blood plasma from a separate cohort of TurboID-transduced and exercised *Albumin-*cre mice. This commercially available anti-CES2 antibody exhibited the expected immunoreactivity to purified, recombinantly produced CES2A and CES2C proteins (**Fig. S3A**). As expected, extracellular CES2 levels from streptavidin-purified *Albumin-*cre secretomes were increased after exercise. (**Fig. 4B**). The total hepatocyte secretome biotinylation signal remained unchanged (**Fig. S3B**), establishing equivalent secretome protein loading. We therefore conclude that CES2 secretion from the liver is a robust molecular event in response to one week of treadmill running.

We sought to test the hypothesis that lactate might serve as an exercise-inducible extracellular signal that stimulates CES2 secretion from hepatocytes. This hypothesis was based on the well-established increase in lactate flux through the liver via the Cori cycle, as well as our previous experiments showing that other metabolic fuels (e.g., fatty acids) can stimulate protein secretion from the liver (Wei et al., 2020). Primary hepatocytes were treated with lactate (2 mM, 4 h) and extracellular CES2 proteins were measured by Western blotting in both cell lysates and conditioned medium. As additional controls, we tested a variety of other exercise-regulated organic acids, including pyruvate, acetate, malate, fumarate, beta-hydroxybutyrate, kynurenate, and pantothenate (Agudelo et al., 2014; Contrepois et al., 2020; Reddy et al., 2020; Sato et al., 2022; Schranner et al., 2020). As shown in **Fig. 4C**, lactate treatment robustly increased the levels of extracellular CES2. Pyruvate, a structurally similar metabolite, also increased CES2 secretion, though with a slightly lower magnitude than that of lactate. By contrast, none of the other metabolites tested increased extracellular CES2 levels (**Fig. 4C** and **Fig. S3C**), establishing that only extracellular lactate, and to a lesser extent pyruvate, exhibit CES2 secretion stimulatory activity.

A dose response of lactate revealed increased CES2 secretion at >2 mM lactate levels (**Fig. 4D**), whereas intracellular CES2 levels were unchanged at all concentrations of lactate tested. In addition, the effect of lactate to induce secretion of CES2 was specific since extracellular albumin levels were unchanged with lactate treatment (**Fig. S3D**). This lactate-induced CES2 secretion is cell type-specific since exogenous expression of CES2A in HEK293T cells resulted in complete retention of this protein intracellularly and treatment of lactate (1-50 mM, 4 h) concentration did not induce the release of CES2A into conditioned medium (**Fig. S3E**). Reduced CES2 secretion was observed when hepatocytes were concurrently treated with lactate (2 mM, 4 h) and an inhibitor of vesicle transport, brefeldin A (BFA 5 µg/ml, 4 h), establishing that these CES2 proteins are released extracellularly via the classical secretory pathway (**Fig. 4E**, and **Fig. S3F**). Lastly, because lactate import into hepatocytes is critical for this metabolite to function as a substrate in the Cori cycle, we next tested whether lactate import via the monocarboxylate transporters (MCTs) was required for induction of CES2 protein secretion. Treatment of primary hepatocytes with AR-C155858, a nanomolar dual MCT1/2 inhibitor (Ovens et al., 2010a; Ovens et al., 2010b), dose-dependently inhibited the lactate-induced secretion of CES2 (**Fig. 4F**). Once again, the inhibitory effect of AR-C155858 was selective for CES2, since no changes were observed in extracellular albumin under these conditions (**Fig. S5G**). These data demonstrate that extracellular lactate is sufficient to drive secretion of CES2 proteins via classical pathway from hepatocytes in a manner that requires import of lactate into hepatocytes (**Fig. 4G**).

### Soluble CES2 proteins exhibit anti-obesity and anti-diabetic effects in mice

Finally, we sought to determine whether release of extracellular CES2 from the liver following exercise was simply a response to exercise training, or whether soluble CES2 proteins might function as circulating molecular effectors of physical activity. Supporting a potential functional role for extracellular CES2, three prior studies showed that liver-specific overexpression of either human or mouse CES2 lowered body weight, reduced hepatic steatosis, and improved glucose homeostasis (Li et al., 2016; Ruby et al., 2017; Xu et al., 2021). An intestine-specific transgenic CES2C mouse model also exhibited a similarly improved metabolic phenotype (Maresch et al., 2019). However, these prior studies did not consider the possibility that extracellular CES2, which is likely also increased in addition to elevation of intracellular CES2 in these transgenic models, might in part mediate the anti-obesity, anti-steatosis, and anti-diabetic phenotypes observed.

To directly test the functional role extracellular CES2 in energy metabolism and glucose homeostasis without a confounding contribution from intracellular CES2, we set out to generate an engineered version of CES2 that would be exclusively localized extracellularly (**Fig. 5A**). Analysis of the primary amino acid sequences for both murine CES2A and CES2C proteins revealed an N-terminal signal peptide, a central alpha/beta hydrolase superfamily domain with the catalytic active site GXSXG motif, and a C-terminal HXEL motif (X=A for CES2A, and R for CES2C). Previous studies showed that the C-terminal HXEL motif is indispensable for the ER lumen localization and C-terminal deleted version of CES2 can be readily detectable in the conditioned medium of cancer cells (Hsieh et al., 2015; Oosterhoff et al., 2005; Potter et al., 1998). We therefore generated CES2A/C constructs in which the HXEL amino acids were removed from the C-terminus (CES2-ΔC) (**Fig. 5A**). An additional N-terminal Flag epitope tag was included after the signal peptide to aid downstream detection. Both wild-type CES2 and CES2-ΔC constructs were transfected into HEK293T cells and the CES2 protein localization was determined by Western blotting of cell lysates and conditioned media. As expected, full-length CES2A/C were enriched intracellularly, whereas both CES2A-ΔC and CES2C-ΔC proteins were exclusively found extracellularly (**Fig. S4A**).

**Fig 5.**
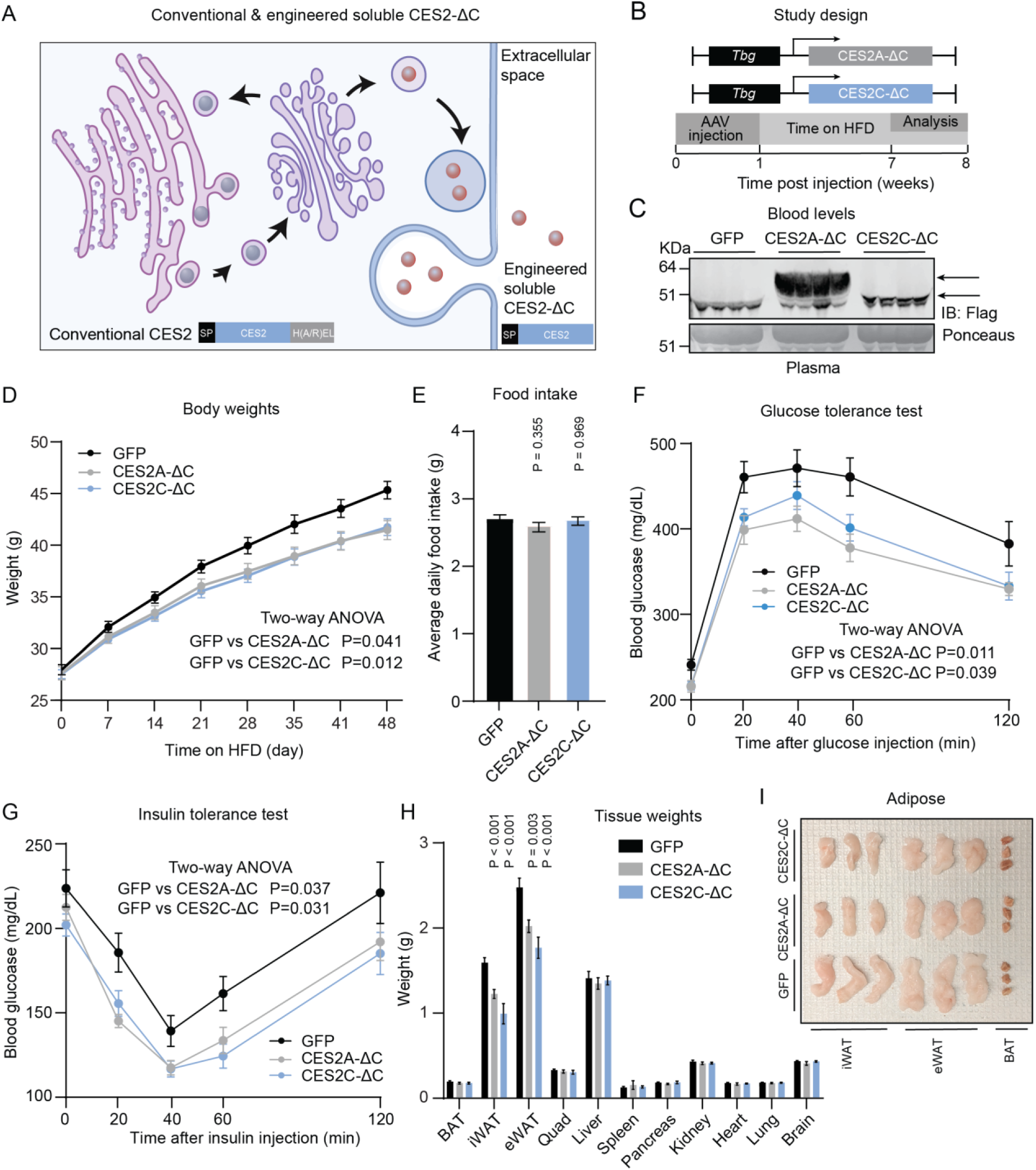
Engineered soluble CES2 have anti-obesity and anti-diabetic effects in mice. (A) Cartoon schematic of conventional ER lumen localized CES2A/C (left) and engineered CES2A/C-ΔC (right). The C-terminal HXEL sequence was removed from conventional CES2A/C to generate engineered soluble CES2A/C-ΔC. (B) Cartoon schematic of the AAV constructs driven by the hepatocyte-specific *Tbg* promoter and study design of HFD feeding experiment. 8 to 10-week-old male C57BL/6 mice were transduced with AAV-*Tbg*-CES2A-ΔC or AAV-*Tbg*-CES2C-ΔC or AAV-*Tbg*-GFP (N = 10/group, 10e11 GC/mouse, intravenously). One-week later, mice were placed with HFD feeding for 7 weeks. In the last week, glucose tolerance test and insulin tolerance test were conducted. At the end of this experiment, tissues and blood were harvested and analyzed. (C) Anti-Flag blotting (top) or loading control (bottom) of blood plasma from 16 to 18-week-old male C57BL/6 mice injected with indicated viruses. N = 4/condition. (D-I) Body weights over the first 7-week of HFD feeding (D) and food intake (measured weekly) (E), glucose tolerance test (F), insulin tolerance test (G), tissue weights (H) and inguinal white adipose tissue (iWAT), epididymal adipose tissue (eWAT) and brown adipose tissue (BAT) after 48 h of 4% PFA fixation (I) from 16 to 18-week-old male C57BL/6 mice injected with indicated viruses for 8 weeks. N = 10/condition. Samples from (I) were from randomly chosen from mice of each treatment group. P-values for (E) and (H) were calculated from two-tailed unpaired t-tests. P-values from (D), (F) and (G) were calculated from two-way ANOVA with post hoc Sidak’s multiple comparisons test.

To deliver the engineered soluble CES2 proteins to mice, we generated adeno-associated virus (serotype 8) expressing each of our two engineered CES2A-ΔC and CES2C-ΔC constructs under the control of the hepatocyte-specific thyroxine binding protein (*Tbg*) promoter (**Fig. 5B**). Next, mice were transduced with AAV-*Tbg*-CES2A-ΔC or AAV-*Tbg*-CES2C-ΔC (N = 10/group, 10e11 GC/mouse, intravenously). Control mice were transduced with an equal titer of AAV-*Tbg*-GFP. As expected, Western blotting of blood plasma using an anti-Flag antibody revealed elevation of circulating CES2A-ΔC and CES2C-ΔC (**Fig. 5C**), which was further validated by measuring plasma ester hydrolysis activity using a previously reported synthetic substrate of carboxylesterases (**Fig. S4B**). In contrast to blood plasma, we did not observe significant changes in total liver CES2 protein level as shown by the anti-CES2 antibody staining (**Fig. S4C**) nor an increase in liver ester hydrolysis activity (**Fig. S4D**). These data confirm that our viral constructs increase only extracellular CES2A and CES2C levels without affecting the intracellular levels of CES2.

One week after viral transduction, mice were placed on high-fat diet (HFD, 60% kcal from fat). Over the subsequent 7 weeks, both CES2A-ΔC and CES2C-ΔC groups of mice exhibited reduced body weight compared to mice transduced with AAV-*Tbg*-GFP (GFP: 45.4 ± 0.8 g versus CES2A-ΔC 41.5 ± 0.9 g and CES2C-ΔC 41.8 ± 0.8 g, mean ± SEM) (**Fig. 5D**). Food intake over this time period was unaltered, suggesting that the lower body weights are not simply due to reduced caloric intake (**Fig. 5E**). In the 7th week, glucose and insulin tolerance tests revealed improved glucose clearance and insulin sensitivity in both CES2A-ΔC and CES2C-ΔC groups (**Fig. 5F** and **G**). Dissection of tissues at the end of the experiment revealed significant reductions of inguinal white adipose tissue (iWAT) (23% and 38% reduction for CES2A-ΔC and CES2C-ΔC, respectively, vs GFP) and epididymal adipose tissue (eWAT) mass (18% and 29% reduction for CES2A-ΔC and CES2C-ΔC, respectively, vs) (**Fig. 5H** and **I**). The lean mass of all three groups remained unchanged (**Fig. 5H**), establishing the effects on body weight are due to reduced adiposity and not any changes in lean mass. Taken together, we conclude that extracellular CES2 proteins have functions in energy balance that are independent of their intracellular roles in triglyceride hydrolysis.

## DISCUSSION

Here we have generated an organism-wide proteomic dataset of the cell type-specific secretome responses to one week of treadmill running. In contrast to previous efforts that have focused on individual secreted factors or individual cell types, our current dataset provides several insights of potential importance to our understanding of cell and tissue crosstalk during physical activity, including 1) the demonstration that ∼20% of cell type-protein pairs exhibit complex, bidirectional, and cell type-specific regulation following exercise; 2) the identification of *Pdgfra* cells as a highly responsive cell type to exercise; and 3) discovery of secreted CES2 as extracellular enzymes with anti-obesity and anti-diabetic functions.

Classically, muscle has been studied as a principal source of activity-inducible “myokines” that mediate tissue crosstalk in exercise. More recent evidence has expanded this model to include exercise-inducible hepatokines from the liver (De Nardo et al., 2022) and adipokines from the fat (Takahashi et al., 2019). In general, many of these past studies have focused on individual cell types. Our studies suggest that many more cell types respond to exercise than previously recognized, including *Pdgfra*+ cells distributed to multiple organ systems. Recent reports using single cell RNA-sequencing showed that exercise regulation in adipose was most strongly pronounced in adipose stem cells (Yang et al., 2021), which are defined by the expression of *Pdgfra*+ (Shin et al., 2020). These adipose-resident fibroblasts may indeed correspond to a subset of the *Pdgfra*+ cells identified in our secretome profling dataset. In the future, it will be important to specifically determine which exercise-regulated cell types and in which organs beyond adipose are defined by *Pdgfra*+ expression.

Using cell culture systems, we also provide evidence that the exercise-inducible secretion of CES2 from the liver can be recapitulated by addition of extracellular lactate to primary hepatocytes in vitro. While the precise downstream mechanism linking lactate import to protein secretion from hepatocytes remains unknown, we suspect one likely possibility includes lactate-inducible proteolytic cleavage of the CES2 C-terminus. This C-terminus contains the ER retention signal required for intracellular localization of CES2. In addition, RHBDL4 has been proposed as a hydrolase that liberates an array of ER-resident proteins into the extracellular space (Tang et al., 2022). Consistent with this idea, we were unable to detect peptides corresponding to C-terminus of either CES2A or CES2C (**Fig. S5**). Such an ER-retention signal cleavage mechanism may also explain the secretion of several other ER-resident proteins in our dataset, such as H6PD (in *Adiponectin-*cre secretomes), CALX (in *Albumin*-cre and *Lysm*-cre secretomes) and AT2A1 (in *MCK*-cre secretomes). The possibility that lactate itself constitutes a more general mechanism linking physical activity to secreted protein-mediated tissue crosstalk remains an open question for future work.

Lastly, we provide evidence that exercise-inducible secretion of CES2 proteins is not simply a molecular response to exercise. Instead, using engineered versions of CES2 that are localized exclusively extracellularly, we show that soluble CES2 proteins exhibit anti-obesity and anti-diabetic actions in obese mice. Interestingly, the metabolic effects of soluble CES2 proteins are not simply due to reduced caloric intake. We hypothesize that CES2 proteins might regulate some other aspect of energy balance, such as energy expenditure or nutrient absorption. Lastly, the *CES2* locus in humans has been linked to multiple cardiometabolic parameters in the UK Biobank, including HDL cholesterol (P = 5.14e-311, beta = +0.0625), blood pressure (P = 1.64e-17, beta = +0.0317), and BMI-adjusted waist-hip ratio (P = 4.61e-15, beta = -0.0297), suggesting that exercise-regulated soluble CES2 proteins might also impact cardiometabolic health in humans.

Projecting forward, an important extension of this work will be to understand the effects of different exercise training regimes and training time on the cellular secretomes in vivo. Which of the secretome responses observed here are specific to the training protocol used in this study, and which might be general responses to multiple types of exercise? Our time course of *Pdgfra*- cre secretomes in response to exercise training already provides evidence that the secretome responses can be complex and time-dependent. Indeed, a single bout versus multiple bouts of either aerobic or resistance training produces distinct phenotypic responses that correlate with activation of overlapping yet distinct molecular pathways (Kraemer and Ratamess, 2005; Wilkinson et al., 2008). On the other hand, our identification of a functional coupling between lactate and CES2 secretion from the liver suggests that soluble CES2 may be a molecular effector more generally associated with multiple and distinct modalities of physical activity. In addition, these murine datasets will need to be compared with studies of exercise plasma in large and deeply phenotyped human cohorts (Robbins et al., 2021; Sanford et al., 2020) to determine which molecular changes might be conserved or species-specific. Ultimately, we anticipate that these and other organism-wide datasets will provide a systematic foundation for probing the role of secreted proteins as mediators of tissue crosstalk in exercise and as molecular effectors of physical activity across the diverse cell and organs systems across the body.

## ACKNOWLEDGEMENTS

We thank members of the Long, Bertozzi and Svensson labs for helpful discussions. We gratefully acknowledge the staff at the Penn Vector Core for the production of AAVs. This work was supported by the US National Institutes of Health (DK105203 and DK124265 to J.Z.L, DK125260 and DK111916 to K.J.S, K99GM147304 to N.M.R.), the Wu Tsai Human Performance Alliance (research grant to J.Z.L, postdoctoral fellowship to X.C.), the Stanford Diabetes Research Center (P30DK116074), the Stanford Cardiovascular institute (CVI), the Jacob Churg Foundation, the McCormick and Gabilan Award (K.J.S), Vanessa Kong-Kerzner foundation (graduate fellowship to W.W.), the American Heart Association (postdoctoral fellowship to M.Z.), and Fundacion Alfonso Martin Escudero (postdoctoral fellowship to M.D.M.G.)

## AUTHOR CONTRIBUTIONS

Conceptualization, W.W., N.M.R., and J.Z.L.; methodology, W.W., N.M.R., and X.L.; investigation, W.W., N.M.R., X.L., X.S., J.G., M.Z., M.D.M.G., H.B., A.T., V.L.L., and W.H.; writing – original draft, W.W., N.M.R., and J.Z.L.; writing – review & editing, W.W., N.M.R., X.L., and J.Z.L.; resources, J.Z.L., C.R.B., K.J.Z., and M.P.S.; supervision and funding acquisition, J.Z.L. and C.R.B.

## STAR METHODS

### Key resources table

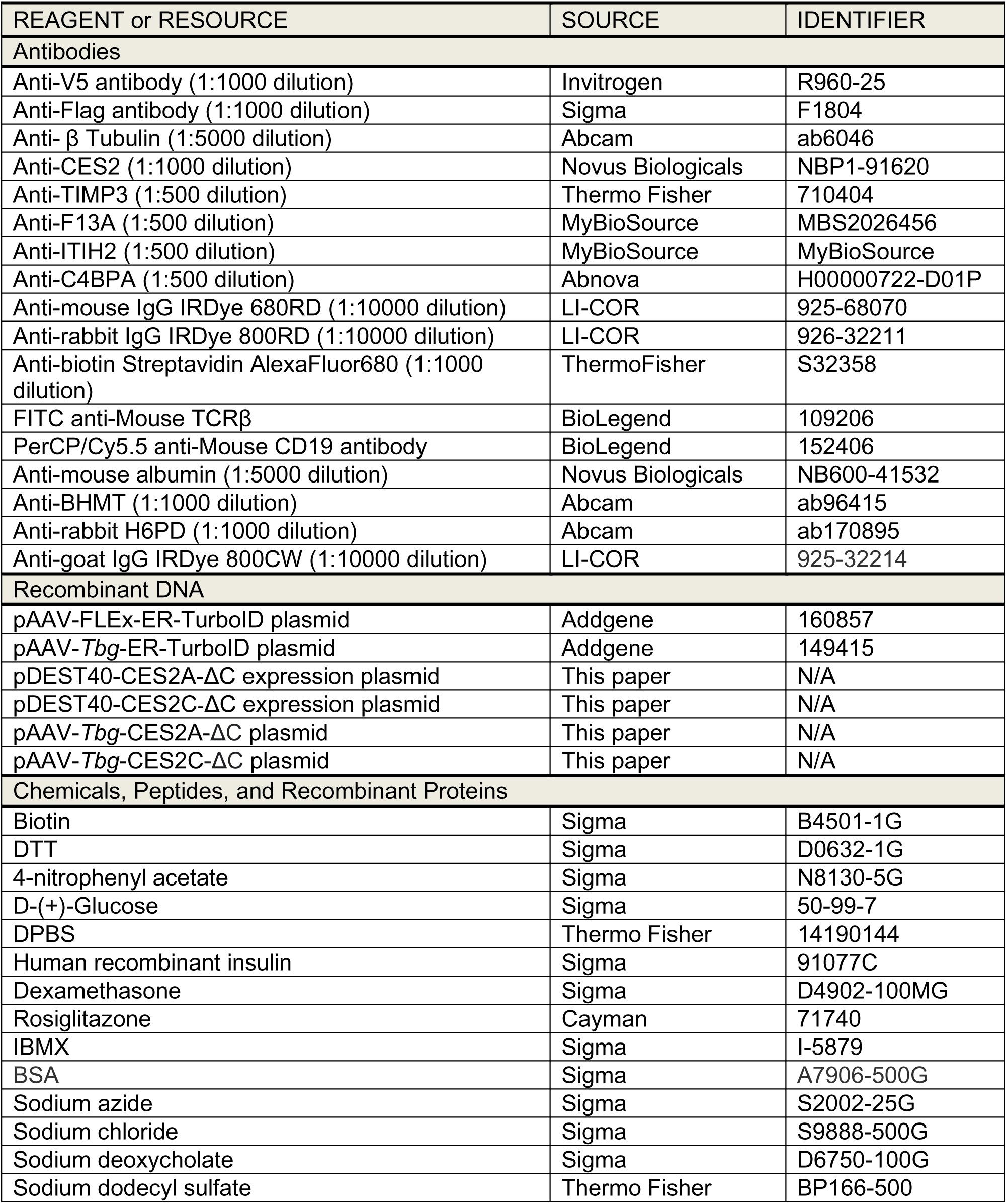
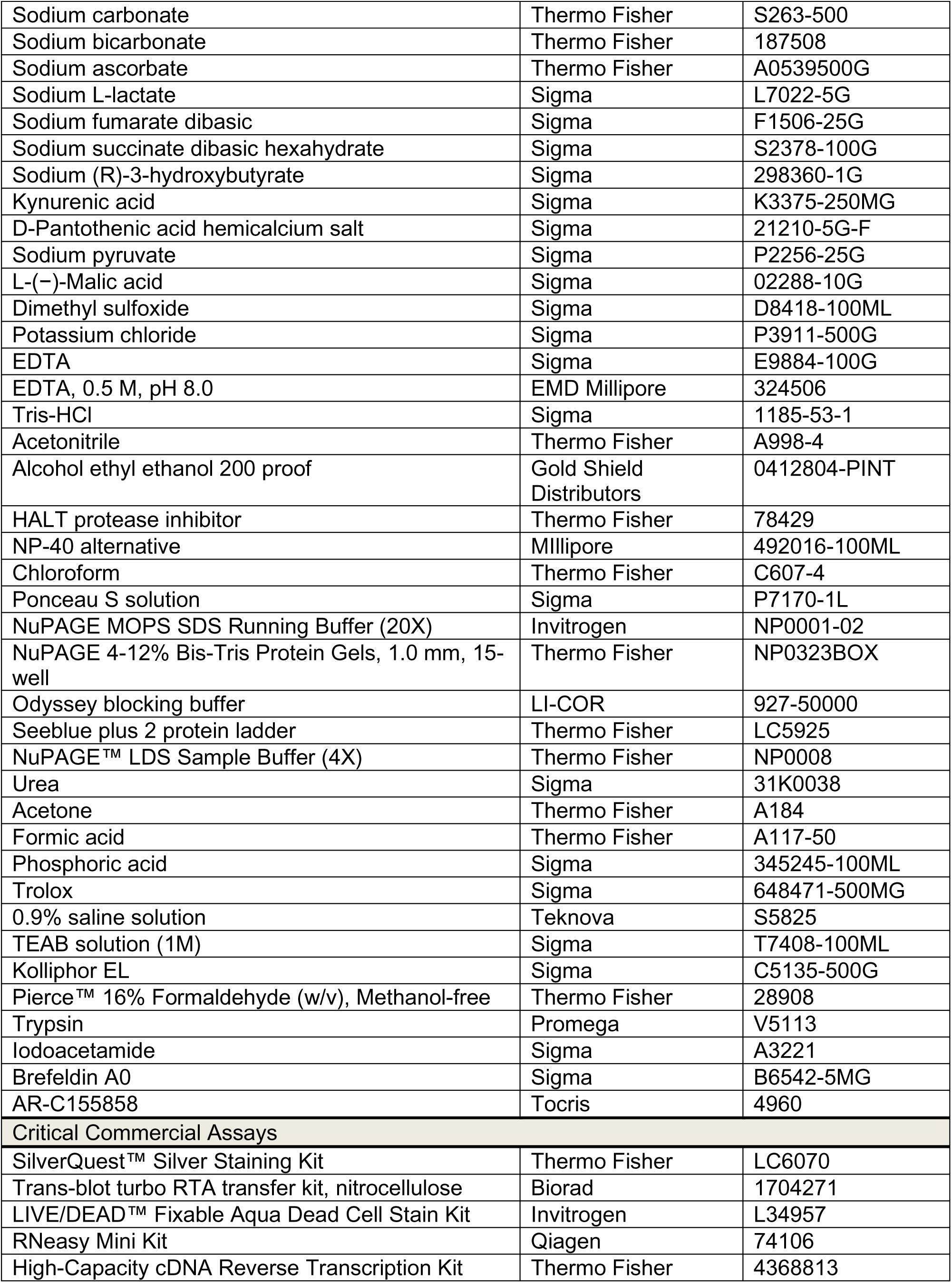
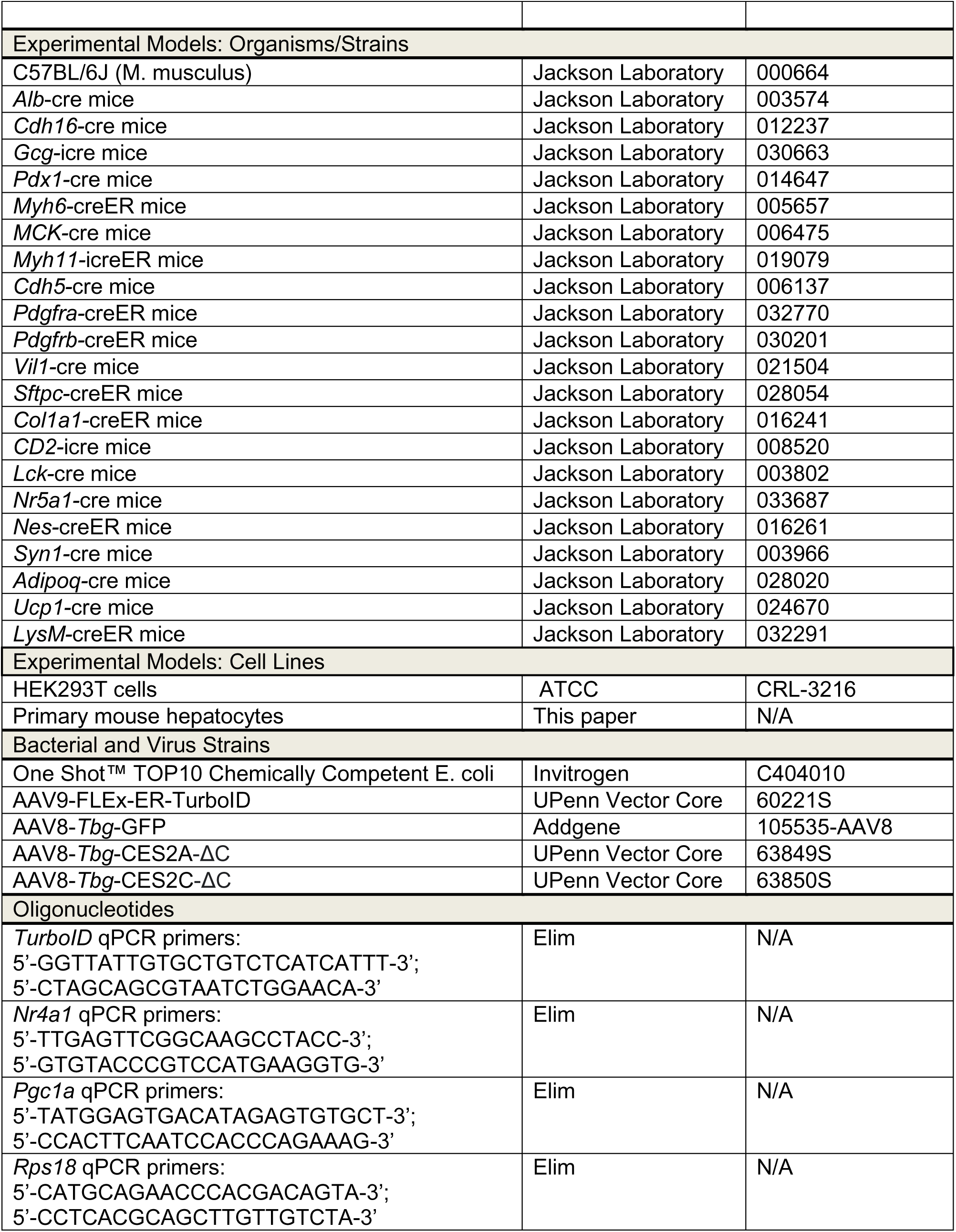
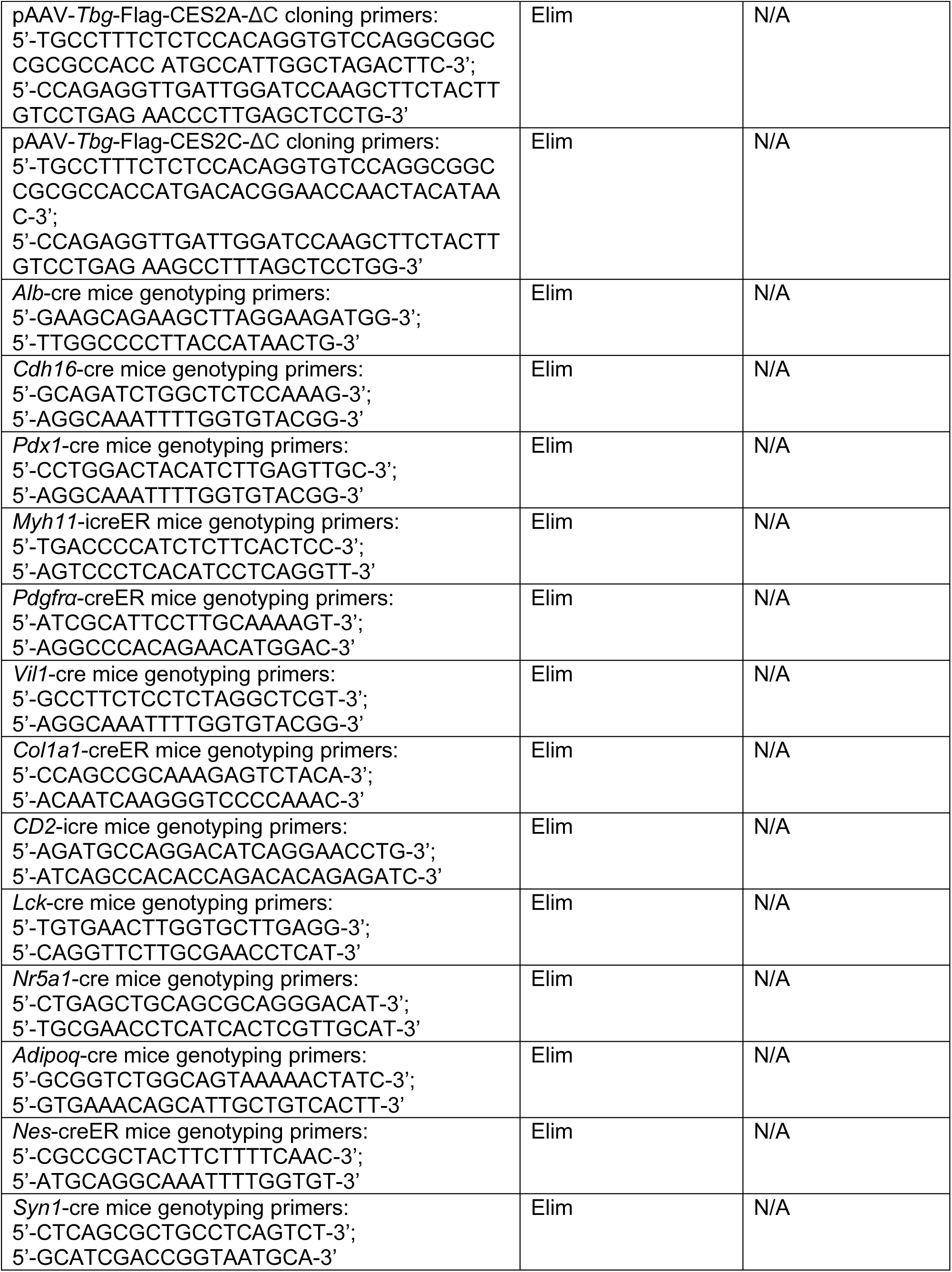
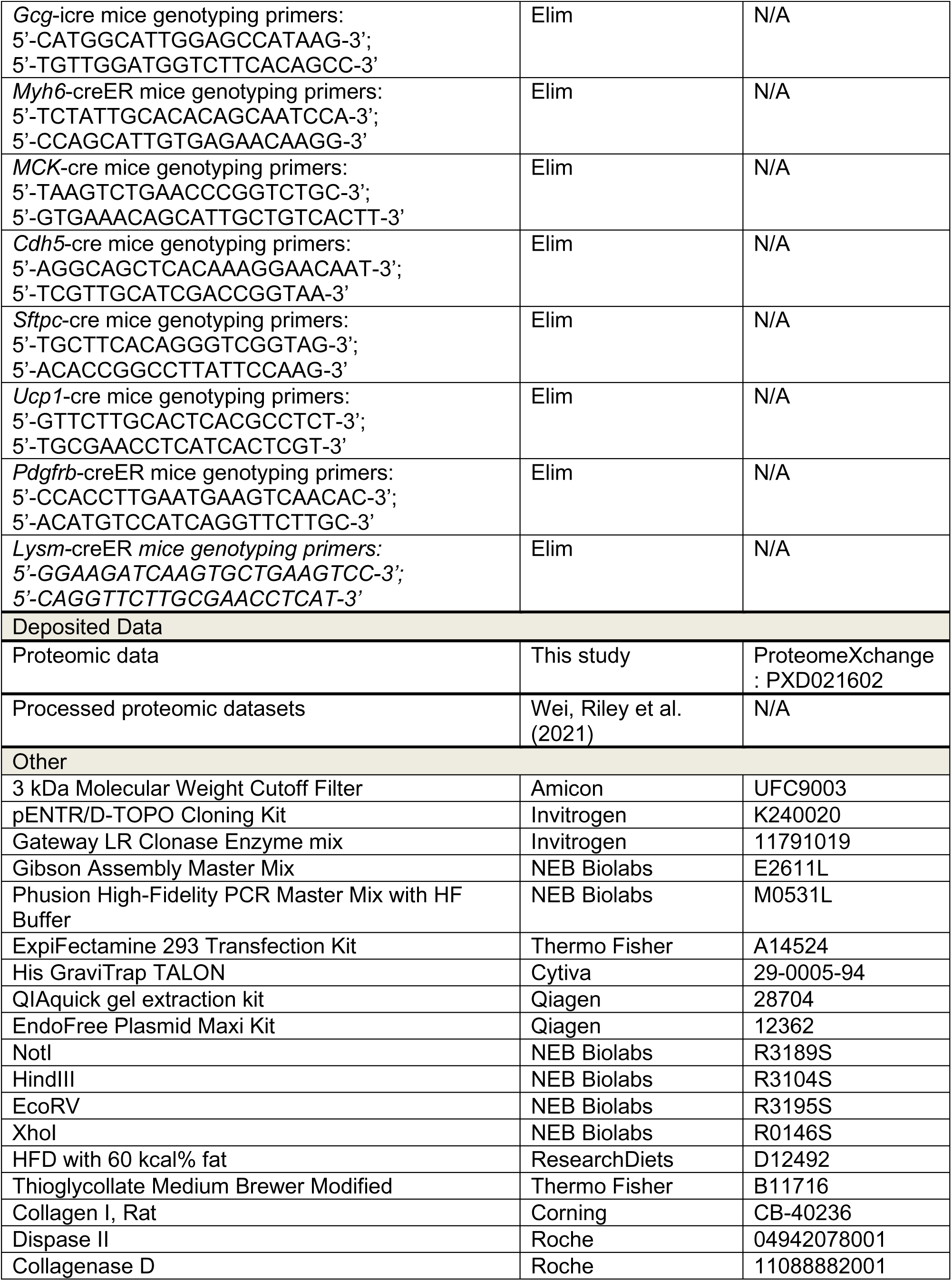
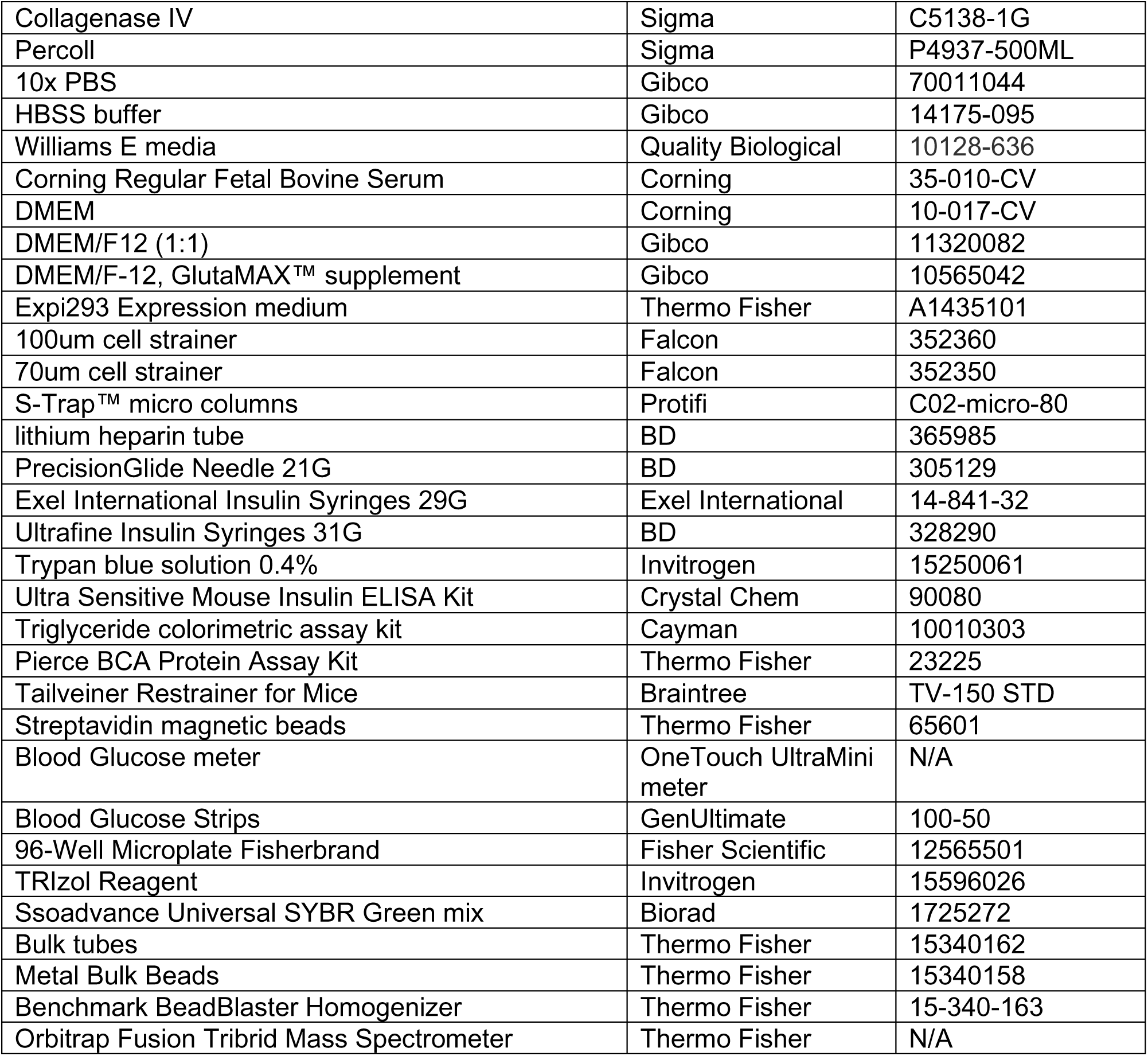

## RESOURCE AVAILABILITY

### Lead contact

Further information and requests for reagents may be directed to and will be fulfilled by the lead contact Dr. Jonathan Z. Long.

### Materials availability

Further information and requests for reagents will be fulfilled by Dr. Jonathan Z. Long. A list of critical reagents (key resources) is included in the key resources table. Material that can be shared will be released via a Material Transfer Agreement.

### Data and code availability

The mass spectrometry proteomics data have been deposited to the ProteomeXchange Consortium via the PRIDE partner repository with the dataset identifier PXD034535 (Perez-Riverol et al., 2022). Processed data from these outputs is available as Supplemental Data files. Any additional information required to reanalyze the data reported in this paper is available from the lead contact upon request.

## EXPERIMENTAL MODELS AND SUBJECT DETAILS

### Mouse models

Animal experiments were performed according to procedures approved by the Stanford University IACUC. Mice were maintained in 12 h light-dark cycles at 22 °C and ∼50% relative humidity and fed a standard irradiated rodent chow diet. Where indicated, high fat diet (Research Diets, D12492) was used. C57BL/6J male and female mice (stock no. 000664), homozygous *Alb*-cre male mice (stock no. 003574), hemizygous *Cdh16*-cre male mice (stock no. 012237), homozygous *Gcg*-icre mice male mice (stock no. 030663), hemizygous *Pdx1*-cre male mice (stock no. 014647), homozygous *Myh6*-creER male mice (stock no. 005657), hemizygous *MCK-*cre male mice (stock no. 006475), hemizygous *Myh11-*icreER male mice (stock no. 019079), homozygous *Cdh5*-cre male mice (stock no. 006137), heterozygous *Pdgfra*-creER male mice (stock no. 032770), homozygous *Pdgfrb*-creER male mice (stock no. 030201), hemizygous *Vil1*-cre male mice (stock no. 021504), homozygous *Sftpc*-creER male mice (stock no. 028054), hemizygous *Col1a1*-creER male mice (stock no. 016241), hemizygous *CD2*-cre male mice (stock no. 008520), hemizygous *Lck*-cre male mice (stock no. 003802), hemizygous *Nr5a1*-creER male mice (stock no. 033687), hemizygous *Nes*-creER male mice (stock no. 016261), hemizygous *Syn1*-cre male mice (stock no. 003966), hemizygous *Adipoq*-cre male mice (stock no. 028020), hemizygous *Ucp1*-cre male mice (stock no. 024670), and hemizygous *LysM* -cre male mice (stock no. 031674) were purchased from Jackson Laboratory. All the cre driver male mice were crossed with female C57BL/6J mice to generate hemizygous/heterozygous mice. Genotypes were verified following genotyping protocols and using the primers listed on the Jackson Laboratory website.

## METHODS DETAILS

### Cell line cultures

HEK293T cells were obtained from ATCC (CRL-3216) and cultured in complete medium (Dulbecco’s Modified Eagle’s Medium, Corning, 10013CV; 10% FBS, Corning, 35010CV; 1:1,000 penicillin–streptomycin, Gibco, 15140-122). Cells were grown at 37 °C with 5% CO_2_. For transient transfection, cells were transfected in 10 cm^2^ at ∼60% confluency using PolyFect (Qiagen, 301107) and washed with complete medium 6 h later.

### Western blotting

For analyzing samples using Western blot, proteins were separated on NuPAGE 4–12% Bis-Tris gels and transferred to nitrocellulose membranes. Equal loading was ensured by staining blots with Ponceau S solution. Blots were then incubated with Odyssey blocking buffer for 30 min at room temperature and incubated with primary antibodies (1:1000 dilution mouse anti-V5 antibody (Invitrogen, R960-25), 1:1000 dilution mouse anti-FLAG antibody (Sigma, F1804), 1:5000 dilution rabbit anti-β-tubulin antibody (Abcam, ab6046), 1:1000 dilution rabbit anti-CES2 antibody (Novus Biologicals, NBP1-91620), 1:500 rabbit anti-TIMP3 antibody (Thermo Fisher, 710404), 1:500 rabbit anti-F13A antibody (MyBioSource, MBS2026456), 1:500 rabbit anti-ITIH2 (MyBioSource, MBS9612213), 1:500 dilution rabbit anti-C4BPA (Abnova, H00000722-D01P), 1:5000 dilution goat anti-albumin antibody (Novus biological, NB600-41532), 1:1000 dilution rabbit anti-H6PD antibody (Abcam, ab170895), 1:1000 dilution rabbit anti-BHMT (Abcam, ab96415), 1:1000 dilution streptavidin Alexa Fluor 680 (Thermo Fisher, S32358)) in blocking buffer overnight at 4 °C. Blots were washed three times with PBST (0.05% Tween-20 in PBS) and stained with species-matched secondary antibodies (1:10000 dilution goat anti-mouse IRDye 680RD (LI-COR, 925-68070), 1:10000 dilution goat anti-rabbit IRDye 800RD (LI-COR, 925-68070), 1:10000 dilution donkey anti-goat IRDye 800CW (LI-COR, 925-32214)) at room temperature for 1 h. Blots were further washed three times with PBST and imaged with the Odyssey CLx Imaging System. Secondary antibodies were not required for imaging blots incubated with streptavidin Alexa Fluor 680 primary antibody.

### AAV production

pAAV-FLEx-ER-TurboID (Addgene, 160857), pAAV-*Tbg*-CES2A-ΔC, pAAV-*Tbg*-CES2C-ΔC plasmids were amplified, extracted using an endotoxin-free Qiagen Maxiprep kit (Qiagen, 12362) and sequence verified. AAV9-FLEx-ER-TurboID (60221S), AAV8-*Tbg*-CES2A-ΔC (63849S), AAV8-*Tbg*-CES2C-ΔC (63850S) viruses were made with Penn Vector Core.

### Viral transduction

For transduction of brain-specific cre driver lines (*Syn1*-cre and *Nes*-creER mice), injection was carried out as previously reported (Gombash Lampe et al., 2014). Briefly, postnatal day 1 pups were anesthetized on ice for 30-60 s. The temporal vein was identified under a dissection microscope and injected with 10e11 genome copies (GC) of AAV9-FLEx-ER-TurboID virus per mouse diluted in a total volume of 30 μl saline (containing 0.3 μl 0.4% Trypan blue solution) with a 31G syringe (BD, 328290). Injected pups were then recovered in hands for 30 s and returned to home cages. For *Syn1*-cre mice, 8 to 9 weeks after injection, 1-week treadmill running was performed as described below on male mice with the correct genotype. For *Nes*-creER mice, 5 to 6 weeks after injection. Tamoxifen (Sigma, T5648-1G) was prepared as a 20 mg/ml solution in corn oil and administered daily for 5 d (100 μl per day, intraperitoneally) to induce recombination. 3 weeks after the final tamoxifen injection, 1-week treadmill running was performed on male mice with the correct genotype. For transduction of cre/icre driver mice (except *Syn1*-cre mice), 6-week-old male hemizygous mice were injected via tail vein with a 29G syringe (Thermo Fisher, 14-841-32) at a dose of 3*10e11 GC per mouse diluted in saline in a total volume of 100 μl per mouse. Three weeks after transduction with the AAV9-FLEx-ER-TurboID virus, 1-week treadmill running was performed. For transduction of creER/icreER mice (except Nes-creER mice), AAV9-FLEx-ER-TurboID virus was injected via tail vein into 6-week-old hemizygous/heterozygous male mice at a dose of 3*10e11 GC per mouse. After a 2-week transduction period, tamoxifen (Sigma, T5648-1G) was prepared as a 20 mg/ml solution in corn oil and administered daily for 5 d (100 μl per day, intraperitoneally) to induce recombination. After the final tamoxifen injection, mice were housed in their home cages for 3 additional weeks before performing 1-week treadmill running. For transduction of C57BL/6J mice, AAV8-*Tbg*-CES2A-ΔC/CES2C-ΔC/GFP viruses were injected via tail vein into 8 to 10-week-old male mice at a dose of 10e11 GC per mouse. One week after viral transduction, mice were fed with HFD (60% fat, Research Diets, D12492). Body weights and food intake were measured every week. After 6 weeks of HFD feeding, glucose tolerance and insulin tolerance tests were performed. At the end of 7 weeks of HFD feeding, tissues and blood were collected for further analysis.

### Mouse exercise and secretome labeling protocols

A 6-lane animal treadmill (Columbus Instruments, 1055-SRM-D65) was used for mouse running. Prior to treadmill running, the bodyweight of individual mice was measured with a tabletop scale. Mice were then acclimated to the treadmill for 5 minutes before running at a speed of 5 m/min for 5 min. Then the speed was increased to 20 m/min and kept constant for 60 min. A serological pipette was used to manually stir the mice to avoid excessive electrical shock during the whole running period. After exercise, mice were returned to home cages. Running was performed for 7 consecutive days in the morning (between 9-12 am). On the fourth day of one-week treadmill running, biotin water (0.5 mg/ml) was supplemented to initiate labeling and kept accessible to mice until the end of the experiment. On the next day, an additional dose of biotin was administered by injection (24 mg/ml, intraperitoneally, in a solution of 18:1:1 saline:Kolliphor EL:DMSO, final volume of 200 µl per mouse per day) 1 h prior to the running for 3 consecutive days.

### Quantitative PCR

We collected the following tissues from the indicated genotypes: liver from *Alb*-cre mice, heart from *Myh6*-creER and *Myh11*-icreER mice, brain from *Syn1*-cre and *Nes*-creER mice, iWAT from *Adipoq*-cre mice, BAT from *Ucp1*-cre mice, lung from *Sftpc*-creER, *Pdgfrα*-creER, *Pdgfrb*-creER and *Cdh5*-cre mice, quadricep muscle from *MCK*-cre mice, intestines from *Vil1*-cre mice, pancreas from *Gcg*-icre and *Pdx1*-cre mice, kidney from *Cdh16*-cre mice, adrenal gland from *Nr5a1*-cre mice, hind limb with muscles removed from *Col1a1*-creER mice. For *Lck*-cre and *CD2*-icre mice, splenocytes were collected by passing the spleen through a cell strainer (Corning, 352350) and resuspended in 2% BSA solution. Splenocytes were stained with FITC anti-mouse TCRβ (BioLegend, 109206), Percp/Cy5.5 anti-mouse CD19 (BioLegend, 152406) and LIVE/DEAD Aqua (Invitrogen, L34957). αβT cells from *Lck*-cre mice were gated on Aqua-CD19-TCRβ+, isolated with FACS, and spun down at 300 g for 5 min at 4 °C for downstream analysis. For *CD2*-icre mice, αβT cells were gated on Aqua-CD19-TCRβ+ and B cells were gated on Aqua-CD19+TCRβ-. Both αβT cells and B cells were sorted with FACS, mixed, and spun down at 300 g for 5 min at 4 °C for downstream analysis. For *Lysm*-creER mice, a new cohort of 6 to 8-week old mice (N = 3) was transduced with AAV9-FLEx-ER-TurboID virus and was injected with tamoxifen to induce cre recombination the same as described before. 3 weeks after the final tamoxifen injection, 1 ml 3% (w/v) thioglycolate solution (Thermo Fisher, B11716) was intraperitoneally injected. 5 days later, 10 ml ice-cold DPBS (Thermo Fisher, 14190144) was intraperitoneally injected to isolate the accumulated macrophages in the peritoneal cavity. Cells were spun down at 400 g for 10 min at 4 °C for downstream analysis. Isolated cells or 30-50 mg of frozen tissues were added to bulk tubes (Thermo Fisher,15340162) containing metal beads and 1 ml TRIzol Reagent (Invitrogen, 15596026). Tissues were then homogenized using a Benchmark BeadBlaster Homogenizer at 4 °C. The mixture was spun down at 13,000 rpm for 10 min at 4 °C to pellet the insoluble materials. RNA was extracted using a RNeasy Mini Kit (Qiagen, 74106) and reverse-transcribed using a High-Capacity cDNA Reverse Transcription Kit (Thermo Fisher, 4368813). Quantitative PCR was performed using Ssoadvance Universal SYBR Green mix (Biorad, 1725272) with a CFX Opus Real-Time PCR instrument. All values were normalized by the ΔΔCt method to *Rps18*. Primer sequences used are described in Key resources table.

### Plasma and tissue sample preparation from mice

2 h after the final bout of running, blood was collected via submandibular bleeding using a 21G needle (BD, 305129) into lithium heparin tubes (BD, 365985) and immediately spun down at 5,000 rpm for 5 min at 4 °C to retrieve the plasma fractions. All tissues were dissected, weighed on a scale, collected into Eppendorf tubes, and immediately frozen on dry ice and stored at −80 °C. Adipose tissues were collected into 4% paraformaldehyde for histology analysis. For western blot analysis, tissues were mixed with 0.5 ml of cold RIPA buffer and homogenized using a Benchmark BeadBlaster Homogenizer at 4 °C. The mixture was spun down at 13,000 rpm for 10 min at 4 °C to pellet the insoluble materials. The supernatant was quantified using a tabletop Nanodrop One and analyzed by western blot. To remove remaining biotin from blood plasma, 200 μl plasma from a single mouse was added with 15 ml PBS and subsequently concentrated 30-fold using 3 kDa filter tubes (Millipore, UFC900324) by spinning down at 4,000 rpm for 1 h. The flowthrough was discarded, and the dilution and centrifugation steps were repeated until a final solution of 500 μl was retrieved at a 9000-fold final dilution. To enrich biotinylated plasma proteins, 200 μl Dynabeads MyOne Streptavidin T1 magnetic beads (Thermo Fisher, 65602) were washed twice with 1ml washing buffer (50 mM Tris-HCl, 150 mM NaCl, 0.1% SDS, 0.5% sodium deoxycholate, 1% NP-40, 1 mM EDTA, 1× HALT protease inhibitor, 5 mM trolox, 10 mM sodium azide and 10 mM sodium ascorbate) and resuspended in 100 μl washing buffer. The beads were then added to the 500 μl biotin-free plasma solution and incubated at 4 °C overnight with rotation. The beads were subsequently washed twice with 1 ml washing buffer, once with 1 ml 1 M KCl solution, once with 1 ml 0.1 M Na_2_CO_3_ solution, once with 1 ml 2 M urea in 10 mM Tris-HCl (pH 8.0), and twice with 1 ml washing buffer. Eppendorf tubes containing beads were vortexed for 3s between each step to ensure thorough washing. Finally, biotinylated proteins were eluted by boiling at 95 °C for 10 min in 60 μl of 2× sample buffer supplemented with 20 mM DTT and 2 mM biotin. Successful enrichment of biotinylated plasma proteins was validated by running the elution sample on NuPAGE 4–12% Bis-Tris gels followed by silver staining (Thermo Fisher, LC6070) according to the instructions from the manufacturer’s protocol.

### Proteomic sample processing

After cooling down to room temperature for 3 min, boiled streptavidin-purified plasma samples (60 μl) were digested using a Mini S-Trap protocol provided by the manufacturer (Protifi, C02-micro-80). As previously described (Wei et al., 2021), cysteine residues were first alkylated by incubating in 30 mM iodoacetamide (Sigma, A3221) in the dark at room temperature for 30 min. Samples were then acidified with phosphoric acid at a final concentration of 1.2%. 420 μl bind/wash buffer (100 mM tetraethylammonium bromide (TEAB) in 90% methanol) was added to each sample. 150 μl samples were loaded onto micro S-trap columns and spun down at 4000 g for 20 s. The flow-through was discarded, and the centrifugation step was repeated until all the solution passed through the column. Following four washes with 150ul bind/wash buffer, 1 μg trypsin (Promega, V5113) was added to the S-trap and incubated at 47 °C for 90 min. After trypsinization, peptides were washed once with 50 mM TEAB (40 μl), once 0.2% formic acid (40 μl), once with a mixture of 50% acetonitrile and 0.2% formic acid (40 μl) and once of 0.2% formic acid in water (40 μl) by spinning down at 1,000g for 60 s. Eluted fraction from each wash was combined, lyophilized, resuspended in 0.2% formic acid, normalized to concentration using a Nanodrop Spectrophotomerter (Thermo Fisher, absorbance at 205 nm), and analyzed by LC-MS/MS. One microliter of each sample was taken and combined into a pooled sampled that was used to make the chromatogram library.

### Proteomics data acquisition

Proteomics data were acquired using a spectrum-library free DIA approach that relies on gas-phase fractionation (GPF) to generate DIA-only chromatogram libraries (Pino et al., 2020a; Searle et al., 2018). Peptides were separated over a 25 cm Aurora Series Gen2 reverse-phase LC column (75 µm inner diameter packed with 1.6 μm FSC C18 particles, Ion Opticks). The mobile phases (A: water with 0.2% formic acid and B: acetonitrile with 0.2% formic acid) were driven and controlled by a Dionex Ultimate 3000 RPLC nano system (Thermo Fisher). An integrated loading pump was used to load peptides onto a trap column (Acclaim PepMap 100 C18, 5 um particles, 20 mm length, Thermo Fisher) at 5 µl/minute, which was put in line with the analytical column 5.5 minutes into the gradient. The gradient was held at 0% B for the first 6 minutes of the analysis, followed by an increase from 0% to 5% B from 6 to 6.5 minutes, and increase from 5 to 22% B from 6.5 to 66.5 minutes, an increase from 22% to 90% from 66.5 to 71 minutes, isocratic flow at 90% B from 71 to 75 minutes, and re-equilibration at 0% B for 15 minutes for a total analysis time of 90 minutes per acquisition. Eluted peptides were analyzed on an Orbitrap Fusion Tribrid MS system (Thermo Fisher). Precursors were ionized was ionized with a spray voltage held at +2.2 kV relative to ground, the RF lens was set to 60%, and the inlet capillary temperature was held at 275 °C.

Six chromatogram library files were collected through six repeated injections of the pooled sample only. Here, the instrument was configured to acquire 4 m/z precursor isolation window DIA spectra using a staggered isolation window pattern (Amodei et al., 2019) from narrow mass ranges using window placements optimized by Skyline. DIA MS/MS spectra were acquired with an AGC target of 400,000 charges, a maximum injection time of 54 ms, beam-type collisional dissociation (i.e., HCD) with a normalized collision energy of 33, and a resolution of 30,000 at 200 m/z using the Orbitrap as a mass analyzer. The six gas-phase fractionation chromatogram libraries were collected with nominal mass ranges of 400-500 m/z, 500-600 m/z, 600-700 m/z, 700-800 m/z, 800-900 m/z, and 900-1000 m/z. The exact windowing scheme was downloaded from https://bitbucket.org/searleb/encyclopedia/wiki/Home (Pino et al., 2020a) and is available in Supplementary Information here. Precursor MS1 spectra were interspersed every 25 scans with an AGC target of 400,000 charges, a maximum injection time of 55 ms, a resolution of 60,000 at 200 m/z using the Orbitrap as a mass analyzer, and a scan range of either 395-505 m/z, 495-605 m/z, 595-705 m/z, 695-805 m/z, 795-905 m/z, or 895-1005 m/z.

For quantitative samples (i.e., the non-pooled samples) the instrument was configured to acquire 25 x 16 m/z precursor isolation window DIA spectra covering 385-1015 m/z using a staggered isolation window pattern with window placements optimized by Skyline (windowing scheme downloaded from the same link as above and available as Supplementary Information here). DIA spectra were acquired with the same MS/MS settings described above. Precursor MS1 spectra were interspersed every 38 scans with a scan range of 385-1015 m/z, an AGC target of 400,000 charges, a maximum injection time of 55 ms, and a resolution of 60,000 at 200 m/z using the Orbitrap as a mass analyzer. The detailed parameters were recorded in **Table S2**. The mass spectrometry proteomics data have been deposited to the ProteomeXchange Consortium via the PRIDE partner repository with the dataset identifier PXD034535.

### Proteomics data analysis to generate cell type-protein pairs

Staggered DIA spectra were demultiplexed from raw data into mzML files with 10 ppm accuracy using MSConvert (Adusumilli and Mallick, 2017) with settings described in Pino et al (Pino et al., 2020a). Encyclopedia (version 1.12.31) (Searle et al., 2018) was used to search demultiplexed mzML files using an internal PECAN fasta search engine called Walnut (Ting et al., 2017) and a reviewed-plus-isoforms mouse proteome database downloaded February 25, 2022 from Uniprot (Consortium, 2021). Walnut settings were: fixed cysteine carbamidomethylation, full tryptic digestion with up to 2 missed cleavages, HCD (y-only) fragmentation, 10 ppm precursor and fragment mass tolerances, and 5 quantitative ions. The chromatogram library resulting from the Walnut search was then used for Encyclopedia searching, where all similar settings to the Walnut search remained the same, and other settings included a library mass tolerance of 10 ppm, inclusion of both b- and y-type fragment ions, and a minimum number of quantitative ions set at 3. Percolator (version 3.1) was used to filter peptides to a 1% false discovery rate using the target/decoy approach and proteins to a 1% protein-level FDR assuming protein grouping parsimony. Resulting data from EncyclopeDIA were checked in Skyline (Pino et al., 2020b) before further processing with Perseus (Tyanova et al., 2016). Proteins were filtered so that only proteins with 2 or more peptides and those that were detected in all three replicates of at least one condition were retained. Data was converted to cell type-protein pairs, and the median value of the cell type-protein pair intensity was compared to the intensity of that protein detected in WT mice control samples. Keratins were manually removed from our dataset as these proteins are frequently detectable contaminants in mass spectrometry experiments (Mellacheruvu et al., 2013). To remove background labeling contaminants, only cell type-protein pairs that showed a greater than 1.5-fold intensity above the median intensity detected in WT samples were retained (Branon et al., 2018). Then cell type-protein pairs with detected protein intensity across all 6 samples (sedentary and exercise) were included for downstream analysis. Next, cell type-protein pairs with variance > 2 ((log2(maximum intensity) – log2(minimum intensity)) > 2 under either sedentary or exercise conditions) were excluded from further analysis (Bourgon et al., 2010). In total, 1272 cell type-protein pairs passed the above filtering criteria and were considered as bona fide cell type-protein pairs.

### Exercise responsiveness scores calculations

Exercise-regulated cell type-protein pairs (adjusted P-values < 0.05) were used for calculating exercise responsiveness scores. Each cell type-protein pair’s exercise responsiveness was calculated as abs(log_10_(adjusted P-values)) x abs(log_2_(exercise fold change)). Exercise fold change of a cell type-protein pair was defined as median protein abundance across three exercise samples divided by median protein abundance across three sedentary samples of the same genotype. Then the exercise responsiveness scores were calculated by the summarization of the exercise responsiveness of each cell type-protein pair from the same cell type.

### Time-course of *Pdgfra*-creER secretomes

AAV9-FLEx-ER-TurboID virus was injected as previously described into 6-week-old hemizygous male *Pdgfra*-creER mice at a dose of 3*10e11 GC per mouse. After a 2-week transduction period, tamoxifen was delivered to induce cre-mediated expression of ER-TurboID. Three weeks after the final tamoxifen injection, mice were divided into three groups (1-day running, 7-day running and sedentary controls, N = 3/condition). 1 h before running, biotin was administered by injection (24 mg/ml, intraperitoneally, in a solution of 18:1:1 saline:Kolliphor EL:DMSO, final volume of 400 µl per mouse per day). The treadmill running was carried out as previously described. For the 1-day running group, mice were sacrificed, and blood and tissue samples were analyzed 2 h after a single bout of running. For 7-day running group, mice were run for 7 consecutive days and blood and tissues were harvested 2 h after the final bout of running. For the sedentary group, biotin was administered, and blood and tissue samples were collected 4 h after biotin delivery.

### Gene ontology analysis

Proteins with P_adj_ <0.05 from *Pdgfra* secretomes were uploaded to online gene ontology analysis tool http://geneontology.org/ (Ashburner et al., 2000). The enriched biological processes were ranked by gene ratio and P-values.

### Isolation and culture of primary mouse hepatocytes

Primary mouse hepatocytes were isolated and cultured as previously described (Jiang et al., 2021; Wei et al., 2020). Briefly, 8 to 12-week-old male mice (C57BL/6J) were sacrificed and perfused with perfusion buffer (1 g/L glucose, 2.1 g/L sodium bicarbonate, 0.4 g/L potassium chloride and 0.2 g/L EDTA in HBSS buffer) via cannulate vena cava for 5 to 8 min and then with digestion buffer (1 mg/ml collagenase IV (Sigma, C5138-1G) in DMEM/F-12 medium) for 5 to 8 min. The liver was then dissected out, cut into small pieces using a razor blade and passed through a 70-μm cell strainer (BD, 352350) to obtain crude hepatocytes. Cells were then spun down at 50 g for 3 min, resuspended in 10 ml plating medium (10% FBS, 1 μM dexamethasone (Sigma, D4902-100MG), 0.1 μM insulin (Sigma, 91077C), 2 mM sodium pyruvate, 1% penicillin–streptomycin in William’s E medium (Quality Biological, 10128-636)) and spun down again at 50 g for additional 3 min. The pellet was resuspended in 10 ml of a 45% Percoll solution in PBS and spun down at 100 g for 10 min to isolate hepatocytes. The final hepatocyte pellet was resuspended in 10 ml plating medium, spun down again at 50 g for 5 min and resuspended in 1 ml plating medium. Cells were counted and plated in a collagen-coated six-well plate at 2 million cells per well. 4 h later, the plating medium was changed to warm maintenance medium (0.1 μM dexamethasone, 1 nM insulin, 0.2% BSA (Sigma, A7906-500G), 2 mM sodium pyruvate, 1% penicillin–streptomycin), and cells were incubated overnight before further treatment.

### Treatment of hepatocytes with organic compounds, MCT inhibitor and Brefeldin A

24 h after plating, primary hepatocytes were washed twice with warm PBS to remove BSA. Then 2 ml William’s E medium containing the indicated concentrations of sodium lactate (Sigma, 05508-5ML), 2 mM sodium fumarate dibasic (Sigma, F1506-25G), 2 mM sodium succinate dibasic hexahydrate (Sigma, S2378-100G), 2 mM sodium (R)-3-hydroxybutyrate (Sigma, 298360-1G), 2 mM kynurenic acid (Sigma, K3375-250MG), 2 mM D-Pantothenic acid hemicalcium salt (Sigma, 21210-5G-F), 2 mM sodium pyruvate (Sigma, P2256-25G), 2 mM L-(−)-Malic acid (Sigma, 02288-10G) was added. The above organic compounds powder was dissolved in ethyl alcohol 200 proof to make 100 mM master stock and diluted accordingly to reach the indicated concentration in medium. 40 µl ethanol was added as negative control. For MCT inhibitor AR-C155858 (Tocris, 4960) and Brefeldin A (Sigma, B6542-5MG), compound power was dissolved in DMSO to make master stock (100 µM for AR-C155858 and 5 mg/ml for Brefeldin A) and diluted accordingly to reach the indicated concentration in medium containing 2 mM sodium lactate. 4 h later, cells and conditioned medium were harvested and analyzed by western blotting as previously described. For HEK293T cells, cells were washed twice with warm PBS 24 h after transfection and incubated with serum-free medium containing indicated concentration of sodium lactate. 4 h later, cells and conditioned medium were harvested and analyzed by western blotting as described below.

### Construction of plasmids for overexpression of CES2A/C-ΔC

Flag-CES2A-ΔC fragment (ref sequence NM_133960.5) and Flag-CES2C-ΔC fragment (ref sequence NM_145603.2) were synthesized as gBlocks with IDT. For both fragments, 5’- GACTACAAGGATGACGACGATAAGGGGGGCGGT-3’ sequences (encoding Flag tag) were inserted after CES2 sequences encoding secretory signal peptide (1-78 nt). For Flag-CES2A-ΔC fragment, C-terminal 5’-CATGCAGAGCTG-3’ sequences (encoding HAEL as ER retention signal peptide) were deleted. For Flag-CES2C-ΔC fragment, C-terminal 5’- CACAGGGAGCTT-3’ sequences (encoding HREL as ER retention signal peptide) were deleted. Both gene fragments were inserted into D-TOPO vector using pENTR/D-TOPO Cloning Kit (Invitrogen, K240020) and shuttled into pDEST40 mammalian expression vector using Gateway LR Clonase Enzyme mix (Invitrogen, 11791019). The pDEST40 plasmids were then transformed into One Shot TOP10 Chemically Competent E. coli (Invitrogen, C404010), extracted and sequence verified. pAAV-*Tbg*-ER-TurboID plasmid (Addgene, 149415) was cut with restriction enzymes NotI and HindIII to generate the backbone vector. Flag-CES2A-ΔC fragment was amplified using primer sets: 5’- TGCCTTTCTCTCCACAGGTGTCCAGGCGGCCGCGCCACCATGCCATTGGCT AGACTTC-3’, 5’-CCAGAGGTTGATTGGATCCAAGCTTCTACTTGTCCTGAGAACCCTTGAGCTCCTG-3’. Flag-CES2C-ΔC fragment was amplified using primer sets: 5’-TGCCTTTCTCTCCACAGGTGTCCAGGCGGCCGCGCCACCATGACACGGAACCAACTACATAAC-3’; 5’-CCAGAGGT TGATTGGATCCAAGCTTCTACTTGTCCTGAGAAGCCTTTAGCTCCTGG-3’. Both PCR products were purified using QIAquick gel extraction kit (Qiagen, 28704) and ligated with the linearized pAAV-*Tbg* vector using Gibson Assembly Master Mix (NEB Biolabs, E2611L). Ligated plasmids were transformed into One Shot TOP10 Chemically Competent E. coli (Invitrogen, C404010), extracted and sequence verified.

### Generation of recombinant CES2 proteins

Recombinant CES2A and CES2C proteins were generated by transient transfection of pDEST40-CES2A/C-ΔC plasmids in mammalian Expi293 cells following the manufacturer’s instructions. Five to seven days after transfection, conditioned medium was collected, and recombinant proteins were purified using a His GraviTrap TALON column and buffer exchanged to PBS. Protein purity and integrity were analyzed by SDS page. Following purification, recombinant proteins were aliquoted and stored at -80 °C to avoid freeze-thaws.

### Determination of CES2A-ΔC and CES2C-ΔC secretion in cell culture

HEK293T cells were transfected as described above. 30 h after transfection, cells were washed twice with PBS and added with 10 ml serum-free medium. 12 h later, conditioned medium (10 ml) was collected and concentrated 20-fold using 10 kDa filter tubes (Millipore, UFC801024) to 500 µl. Concentrated conditioned medium was mixed with 4× loading buffer (NuPAGE LDS Sample Buffer, Invitrogen, NP0008, 100 mM DTT) and boiled for 10 min at 95 °C. Cells were collected and lysed by probe sonication in RIPA buffer (1% NP-40, 0.1% SDS, 0.5% sodium deoxycholate and 1:100 HALT protease inhibitor, Thermo Fisher, 78429) for 1 min. Cell lysates were spun down at 13000 rpm for 10 min at 4 °C. The supernatant was collected, quantified using a tabletop Nanodrop One, boiled for 10 min at 95 °C. Both conditioned medium and cell lysate samples were then analyzed by western blot.

### Histology

Adipose tissues were collected into 4% paraformaldehyde and fixed at 4 °C with rotation for 72 h. Fixing solution was replaced with 20% sucrose solution. Adipose tissues were dehydrated for additional 24 h before freezing in OCT-embedded block (Thermo Fisher, 23-730-571) and cryosectioned. H&E staining was conducted on the slides by Stanford Animal Histology Core.

### Glucose tolerance and insulin tolerance tests in mice

For glucose tolerance tests, mice were fasted for 6 h (fasting starting 8 am in the morning) and then intraperitoneally injected with glucose at 2 g/kg body weight. Blood glucose levels were measured at 0, 20, 40, 60, and 120 mins via tail bleeding using a glucose meter. For insulin tolerance tests, mice were fasted for 6 h (fasting starting 8 am in the morning) and then intraperitoneally injected with insulin in saline 0.75 U/kg body weight. Blood glucose levels were measured at 0, 20, 40, 60, and 120 mins via tail bleeding using a glucose meter.

### Carboxylesterases enzymatic activity measurement

A continuous spectrophotometric assay was performed using 4-nitrophenyl acetate (Sigma, N8130-5G) as substrate as previously described (Ross and Borazjani, 2007). Briefly, 1 mM 4-nitrophenyl acetate was prepared freshly in 50 mM Tris⋅Cl buffer (pH 7.4). 150 μl of 4-nitrophenyl acetate solution was added into a single well of a 96-well plate (Thermo Fisher, 125565501), followed by pre-incubating 5 min at 37 °C in the absorbance plate reader. 15 μg liver lysates or 3 μl plasma were diluted in 50 mM Tris⋅Cl buffer (pH 7.4) and added to 4-nitrophenyl acetate solution (total volume 300 μl). The formation of p-nitrophenolate was measured every 30 s at 405 nm for 5 min. Subtracted from background absorbance, absorbance at each time point was used to generate a kinetic plot for each sample. Data points within the linear range of the reaction were used to calculate the slope of the enzymatic reaction. Finally, relative enzymatic activity was calculated comparing the slopes of reaction from each sample.

## QUANTIFICATION AND STATISTICAL ANALYSIS

### Data representation and statistical analysis

All values in figures are shown as mean ± SEM. The number of biological replicates (N) is described in each figure legend (N corresponds to the number of animals used under each condition for animal experiments and corresponds to the number of independently conducted experiments for cellular experiments). For animal studies, mice were randomly assigned to control and treatment groups. Each animal study was repeated at least twice using separate cohorts of mice. To identify statistically changed cell type-protein pairs (exercise vs sedentary, N = 3/protein/condition/genotype), protein intensities were first scaled by the scale() function in R package (Alvarez-Castelao et al., 2017). The Limma package (Law et al., 2016; Ritchie et al., 2015) was implemented to conduct the moderated t-statistics (Smyth, 2004) (https://github.com/leolove2022/ModeratedtTest.git), and adjusted P-values of each cell type-protein pair were generated into excel files. 256 cell type-protein pairs out of total 1272 pairs were defined as under exercise regulation (adjusted P-values < 0.05). Each in vitro experiment using primary cells from mice was repeated using at least three cohorts of mice. Two-tailed, unpaired student’s t-test was used for single comparisons assuming the sample groups exhibited a normal distribution and comparable variance. Two-way ANOVA with post hoc Sidak’s multiple comparisons test with repeated measures was used for the body weights, glucose tolerance test and insulin tolerance test studies. Unless otherwise specified, statistical significance was set at adjusted P-value < 0.05 for the proteomics data, and P-value < 0.05 for all other comparisons.

## SUPPLEMENTAL INFORMATION TITLES AND LEGENDS

**Fig S1. Characterization of secretome labeling, running protocol, and secretomes in mice. Related to Figure 1.**

(A) mRNA expression of TurboID in tissues or isolated primary cells, where cre recombinase expression was reported, of wildtype male C57BL/6 mice and 21 male cre driver lines (N = 3/genotype) after AAV9-FLEx-ER-TurboID viral transduction (3*10e11 GC/mouse, intravenously) (see **Methods**).

(B) Anti-V5 and anti-tubulin (loading control) blotting of murine tissues from wildtype mice C57BL/6 or indicated cre driver lines following transduction of AAV9-FLEx-ER-TurboID virus. This experiment was repeated with three biological replicates per genotype and similar results were obtained.

(C-E) mRNA expression of indicated genes in quadricep muscles (C), tissue mass (D) and H&E staining (E) of inguinal white adipose (iWAT) from 8 to 12-week-old male C57BL/6 mice harvested two hours after the final bout of 1-week treadmill running (red circles) or without exercise training (blue circles). N = 5/condition. This experiment was repeated twice, and similar results were obtained.

(F) Body weight change of wildtype male C57BL/6 mice and 21 male cre driver lines over 1-week treadmill running or without exercise training over 1-week period (N = 3/genotype/condition, in total 132 mice).

**Fig S2. Characterizations of *Pdgfra*+ cells secretomes. Related to Figure 3.**

(A) Relative *TurboID* mRNA levels from indicated tissues of heterozygous *Pdgfra*-cre mice (12-week-old male, N=3) injected with 3*10e11 GC/mouse AAV9-FLEx-ER-TurboID and tamoxifen. Controls samples were from C57BL/6 male mice (10-week-old male, N = 3) injected with 3*10e11 GC/mouse AAV9-FLEx-ER-TurboID.

(B) Mean *Pdgfra* gene expression of indicated cell types from *Tabula Muris*.

**Fig S3. Characterizations of lactate induced CES2 secretion in vitro. Related to Figure 4.**

(A) Anti-CES2 blotting of recombinant CES2A proteins (top left), recombinant CES2C proteins (top right) and ponceaus stain (bottom). 8 μg of recombinant proteins generated from Expi293 cells were loaded into each lane.

(B) Anti-biotin blotting of immune purified biotinylated plasma proteins from 10-week-old Albumin-cre male mice transduced with 3*10e11 vg AAV9-FLEx-ER-TurboID virus and exercised on a treadmill for 1-week. N = 5/group.

(C) Anti-CES2 (second row) blotting of cell lysates, and ponceaus stain of conditioned medium (top row) and cell lysates (bottom row) of conditioned medium of primary hepatocytes isolated from 8 to 12-week-old male C57BL/6J mice. Cells were treated with 2 mM indicated organic compounds for 4 h before analysis. Ponceaus staining was used as loading control. This experiment was repeated four times and similar results were obtained.

(D) Anti-CES2 stain (second row) and ponceaus stain (bottom row) of cell lysates and anti-Albumin stain (top row) of conditioned medium from primary mouse hepatocytes treated with indicated concentration of sodium lactate. Primary hepatocytes were isolated from 8 to 12-week-old male C57BL/6J mice. Conditioned medium was collected 4 h after the addition of sodium lactate. Experiments were repeated four times, and similar results were obtained.

(E) Anti-Flag blotting of cell lysates and conditioned medium of HEK293T cells transfected with Flag-CES2A construct and treated with indicated concentrations of sodium lactate. Ponceaus staining was used as loading control. 36 h after transfection, cells were washed and replaced with serum-free medium containing indicated concentrations of sodium lactate for 4 h. Cells and conditioned medium were then collected and analyzed. Experiments were repeated three times and similar results were obtained.

(F) Anti-CES2 (top row), anti-Albumin (second row) and anti-BHMT (third row) blotting of conditioned medium, and anti-CES2 blotting (fourth row) and ponceaus stain (bottom row) of cell lysates of primary hepatocytes treated with indicated concentrations of sodium lactate and BFA. Primary hepatocytes were isolated from 8 to 12-week-old male C57BL/6J mice. Conditioned medium was collected 4 h after the addition of sodium lactate (2 mM) and BFA (5 μg/ml). This experiment was repeated three times, and similar results were obtained.

(G) Anti-Albumin blotting (top row) of conditioned medium and anti-CES2 blotting (second row) and ponceaus stain (bottom row) of cell lysates from primary mouse hepatocytes treated with indicated concentration of sodium lactate and AR-C155858. Primary hepatocytes were isolated from 8 to 12-week-old male C57BL/6J mice. Conditioned medium was collected 4 h after the addition of sodium lactate and AR-C155858. Experiments were repeated four times, and similar results were obtained.

**Fig S4. Engineered soluble CES2 in cells and in mice. Related to Figure 5.**

(A) Anti-Flag blotting of cell lysates (top left) and conditioned medium (top right) of HEK293T cells transfected with Flag-CES2A, Flag-CES2A-ΔC, Flag-CES2C, Flag-CES2C-ΔC constructs. Ponceaus staining was used as loading control (bottom). 36 h after transfection, cells were washed and replaced with serum-free medium for 12 h. Cells and conditioned medium were then collected and analyzed. Experiments were repeated three times and similar results were obtained. (B, D) Relative ester hydrolysis activities of blood plasma (B) and liver lysates (D) from 16 to 18-week-old male C57BL/6 mice 8 weeks after being transduced with indicated viruses. N = 10/condition.

(C) Anti-CES2 (top) and anti-tubulin (loading control, bottom) blotting of liver lysates from 16 to 18-week-old male C57BL/6 mice 8 weeks after being transduced AAV-*Tbg*-CES2A-ΔC (right) or AAV-*Tbg*-GFP (left). N = 7/condition.

**Fig S5. CES2A and CES2C protein sequences.**

Schematic of detected peptides for CES2A (top) and CES2C (bottom) protein mapped onto its respective reference sequences with annotated domains indicated below. Pink box indicates detected peptides, grey box indicates undetected region, black box indicates signal peptide, blue box indicates ER retention peptide (HAEL for CES2A and HREL for CES2C), and black line indicates potential N-glycosylation site.

**Table S1.** Complete LC–MS/MS proteomic analysis of streptavidin-purified plasma proteins from mice transduced with AAV-FLEX-ER-TurboID viruses or control mice.

**Table S2.** Detailed parameters of proteomics data acquisition using DIA mode.

